# Multiscale networks in multiple sclerosis

**DOI:** 10.1101/2023.02.26.530153

**Authors:** Keith E. Kennedy, Nicole Kerlero de Rosbo, Antonio Uccelli, Maria Cellerino, Federico Ivaldi, Paola Contini, Raffaele De Palma, Hanne F. Harbo, Tone Berge, Steffan D. Bos, Einar A. Høgestøl, Synne Brune-Ingebretse, Sigrid A. de Rodez Benavent, Friedemann Paul, Alexander U. Brandt, Priscilla Bäcker-Koduah, Janina Behrens, Joseph Kuchling, Susana E Asseyer, Michael Scheel, Claudia Chien, Hanna Zimmermann, Seyedamirhosein Motamedi, Josef Kauer-Bonin, Julio Saez-Rodriguez, Melanie Rinas, Leonidas G Alexopoulos, Magi Andorra, Sara Llufriu, Albert Saiz, Yolanda Blanco, Eloy Martinez-Heras, Elisabeth Solana, Irene Pulido-Valdeolivas, Elena Martinez-Lapiscina, Jordi Garcia-Ojalvo, Pablo Villoslada

## Abstract

Complex diseases such as Multiple Sclerosis (MS) cover a wide range of biological scales, from genes and proteins to cells and tissues, up to the full organism. We conducted a multilayer network analysis and deep phenotyping with multi-omics data (genomics, phosphoproteomics and cytomics), brain and retinal imaging, and clinical data, obtained from a multicenter prospective cohort of 328 patients and 90 healthy controls. Multilayer networks were constructed using mutual information, and Boolean simulations identified paths within and among all layers. The path more commonly found from the boolean simulations connects MP2K, with Th17 cells, the retinal nerve fiber layer (RNFL) thickness and the age related MS severity score (ARMSS). Combinations of several proteins (HSPB1, MP2K1, SR6, KS6B1, SRC, MK03, LCK and STAT6)) and immune cells (Th17, Th1 non-classic, CD8, CD8 Treg, CD56 neg, and B memory) were part of the paths explaining the clinical phenotype. Specific paths identified were subsequently analyzed by flow cytometry at the single-cell level.

**Graphical abstract:** 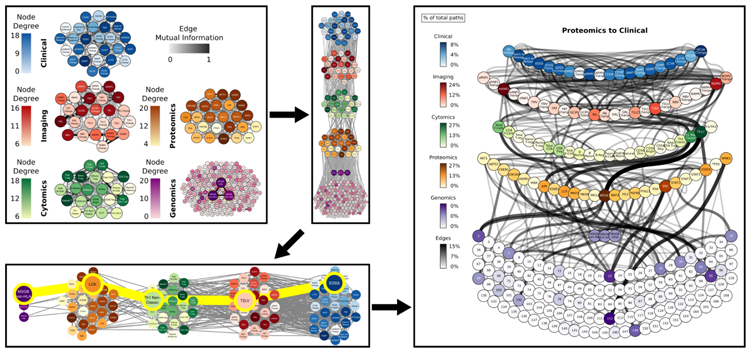

**Author Summary:** Complex diseases such as Multiple Sclerosis (MS) involve the contribution of a wide range of biological processes. We conducted a systems biology study of MS based on network analysis and deep phenotyping in a prospective cohort of patients with clinical, imaging, genetics, and omics assessments. The gene, proteins and cell paths explained variation in central nervous system damage, and in metrics of disease severity. Such multilayer paths explain the different phenotypes of the disease and can be developed as biomarkers of MS.

## Introduction

Complex diseases involve the interaction of multiple biological scales, including tissues, cells, and molecules (genes, proteins, and metabolites), all of which regulate biological function and modulate the susceptibility to a given clinical phenotype. Although significant efforts have been devoted to understanding each of these levels, few attempts have succeeded in integrating multiple scales and the flow of information across them. Such integration would definitely improve our understanding of disease pathogenesis (1, 2) and wellness (3). Multilayer networks provide a framework to integrate complex biological data across different scales, which should allow us to understand the flow of biological information in health and disease (4–6). This is especially important in diseases with a complex genetic and molecular basis, such as Multiple Sclerosis (MS).

MS is an autoimmune disease characterized by inflammatory attacks to the central nervous system (CNS), which damages the neural tissue and leads to significant disability (7). The inflammation occurs in acute attacks as well as by chronic inflammation, defining the different clinical subtypes of the disease, namely relapsing-remitting (RRMS) and progressive (PMS). MS is an example of a complex disease, with different biological scales participating in its pathogenesis, including genetic factors (8), cellular signaling (9, 10), adaptive and innate immunity (11, 12), and CNS damage (13). Additionally, the interplay between these various components is modulated by environmental factors (14, 15), with viral infections and especially the Epstein-Barr virus being the main triggers (16). As a result, the MS phenotype of neurological disability is very heterogeneous and difficult to predict (7, 17), creating significant limitations for patient care. As an example of the difficulty of finding biological determinants of MS, although more than 200 genetic polymorphisms have been associated with MS susceptibility, their contribution to the clinical phenotype is small and remains to be clarified (18). Similarly, many studies have attempted to identify biomarkers of the clinical course and prognosis of the disease, including oligoclonal bands, neurofilament light chain protein, brain or spinal cord volume or retinal thickness, but few have been validated, and even their individual predictive ability is small, making their use in clinical practice limited (19).

Several studies have attempted to integrate biological networks in MS, mainly at the genetic level (20–23). Those studies addressed the biomolecular aspects of the disease (genes and proteins), but they did not describe the relation of those features with tissue damage or clinical disability. In contrast, our approach focuses on bridging the gap between the microscopic and macroscopic scales of MS to better explain the endotype-phenotype relationship. To that end, we use multilayer network analysis to assess how information flows across biological scales, and to identify multiscale paths that contribute to explain the phenotype of MS.

Within the umbrella of the Sys4MS project (24), we recruited a multicenter prospective cohort of 328 patients with MS and 90 healthy subjects with a two-year follow-up and performed deep phenotyping by collecting multi-omics data, imaging, and clinical outcomes. This collection provided data on five biological layers: (1) genes, (2) phosphoproteins (mostly kinases), (3) immune cells, (4) tissue (imaging), and finally (5) the clinical phenotype (**Figure 1a**). Network generation was first applied to each of these layers individually, using mutual information to capture linear and non-linear dependencies between the elements of each layer (**Figure 1b-f**) before the layers were interconnected (**Figure 1g**). Our approach is hypothesis-based, rather than data-based: First, we make use of a set of single nucleotide polymorphisms (SNPs), proteins and immune cell subtypes already known to be associated with MS (8, 9, 24, 25). Second, we consider the transfer of information from genes to proteins and cell layers, which will define the tissue (imaging) and clinical outcomes as the phenotype (**Figure 1g**). In order to obtain functional information from the network models, dynamical simulations using Boolean network modeling were used to identify several paths spanning these five layers.

**Figure 1.**
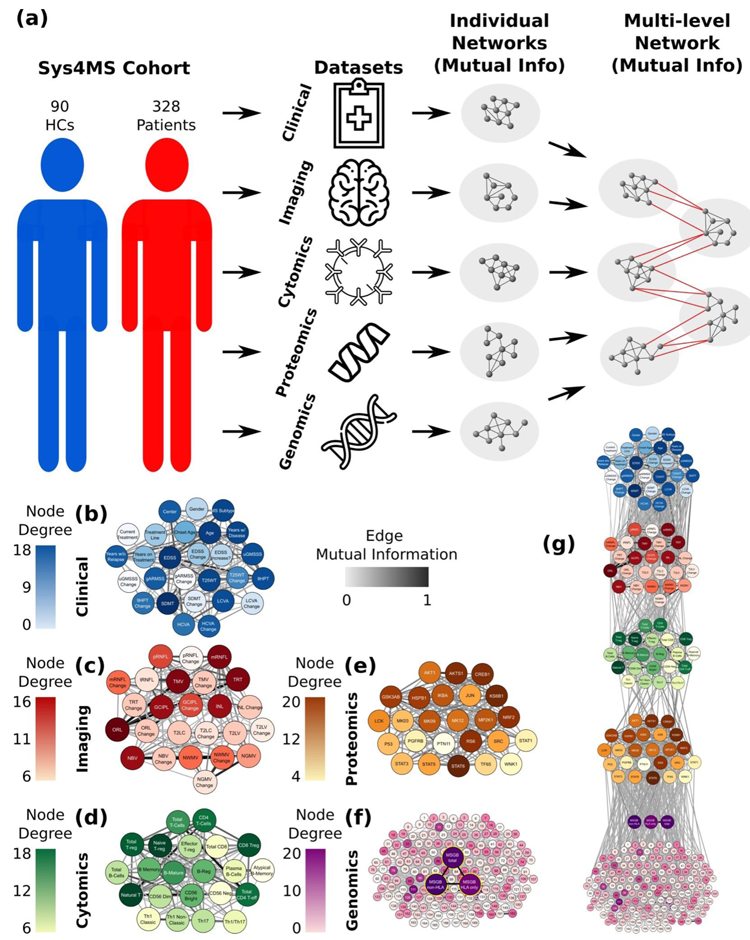
Building multilayer networks using multi-omics, imaging, and clinical data. (a) Illustration of network construction. The data from each layer is taken from the cohorts and used to create networks, where the nodes are the elements in the dataset (genomics, phosphoproteomics, cytomics, tissue imaging, and clinical data), and the edges correspond to the mutual information between element pairs across all subjects. Once individual networks are created, they are linked together, again using mutual information, following a hierarchy that connects each layer successively, starting with genomics and working up to the phenotypic (clinical) layer. (b-f) Topology of individual layer networks from the experimental data. In each of the networks, the degree of each node is color-coded, with higher degrees in darker colors. The edge weights are coded in grey scale in a similar manner, with a darker edge representing a higher weight, and thus a higher correlation between nodes. The genomics network was enriched with the previous knowledge on regulatory networks (f) and included the MS genetic burden scores (g). In the combined five-layer network, the layers are connected using the hierarchy described above, with genomics at the bottom and clinical phenotype at the top. High resolution network representations for single-layer networks are available at Github link https://keithtopher.github.io/single_networks/<u>#/</u> and for multilayer networks at https://keithtopher.github.io/combo_networks/<u>#/</u>.

## Results

The focus of the results is on the paths between the genes, proteins, cells and the phenotype (imaging and clinical scales). Each step below shows how the paths were identified, and which sources tend to be more strongly connected with the phenotype. First, descriptive information about the data is given, then the networks of the layers are constructed, then Boolean simulations are run, and finally the top paths are selected.

### Deep phenotyping: multi-omics, imaging, and clinical data from MS patients

We recruited 328 MS patients (age 41±10 years, 70% female) at four centers throughout Europe, corresponding to the Sys4MS cohort (**Table 1**). Of these, 271 patients (82%) had RRMS, and 57 (17%) had PMS. We also recruited 90 healthy controls (HCs) matched by sex and age with the RRMS group. The patients had a mean disease duration of 10 (SD 8) years, and median Expanded Disability Status Scale (EDSS) of 2.0 (range: 0-8). Regarding the use of disease modifying drugs (DMD) at baseline, 70% of patients were treated, 44% with low-efficacy therapies, and 26% with high-efficacy therapies (see **Methods** for drug definition). By the second year of follow-up (mean follow-up 1.98 ± 0.94 years, n=274), two RRMS cases progressed to PMS, 22 patients started new therapies (cladribine: 1; fingolimod: 2; glatiramer acetate: 4; ocrelizumab: 9; rituximab: 2; teriflunomide: 4) and 17 changed from low to high-efficacy therapies. Imaging data consisted of both brain magnetic resonance imaging (MRI) and retina optical coherence tomography (OCT) **(Table 1)**.

**Table 1.**
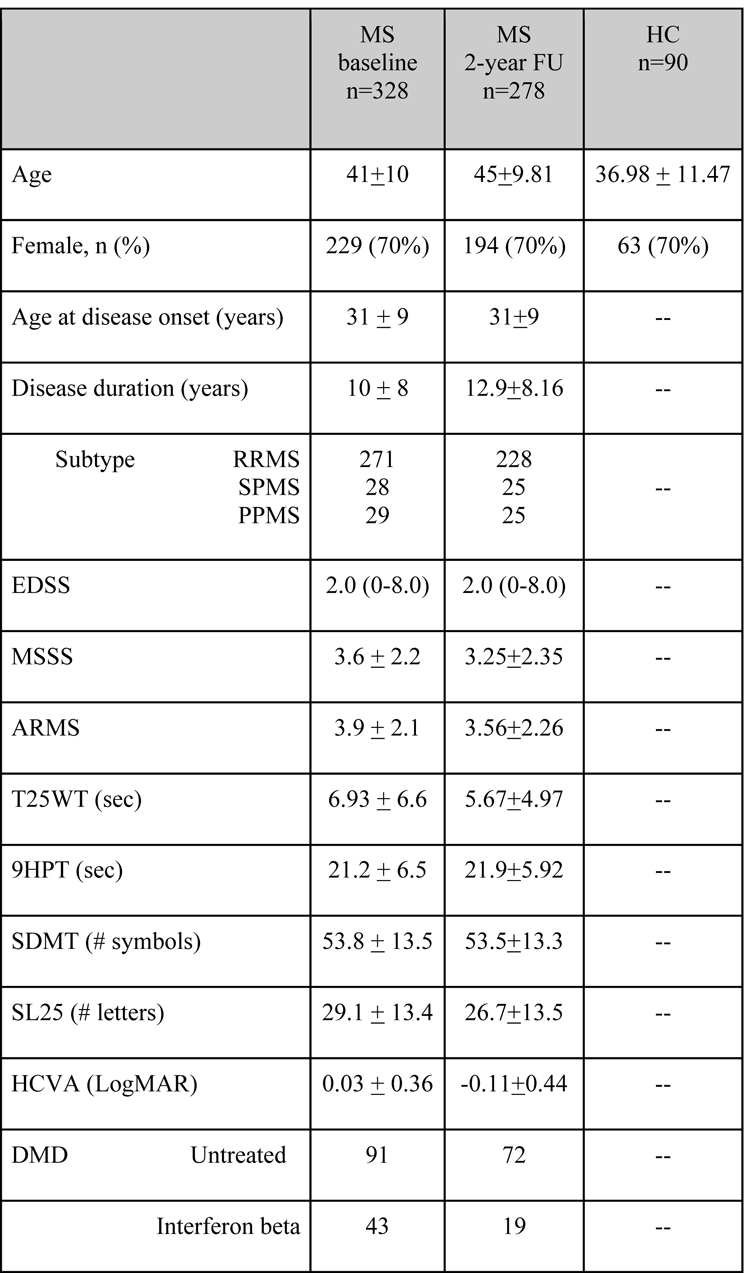
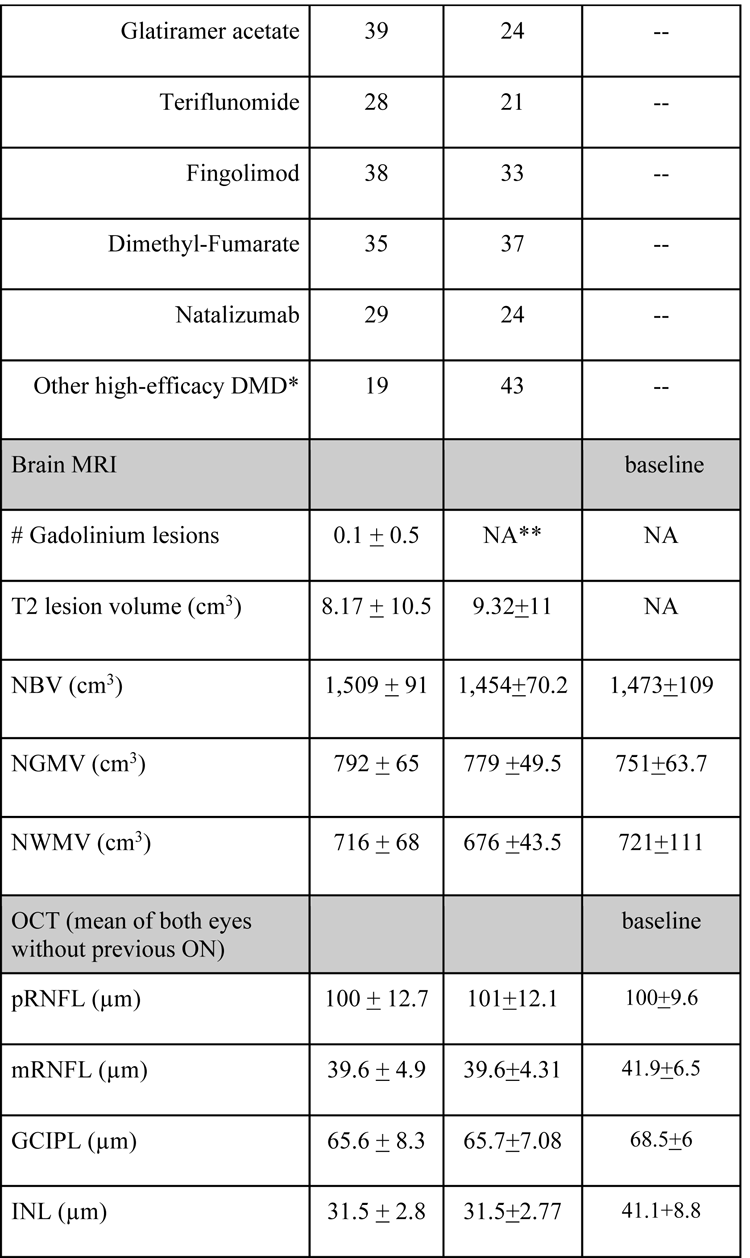
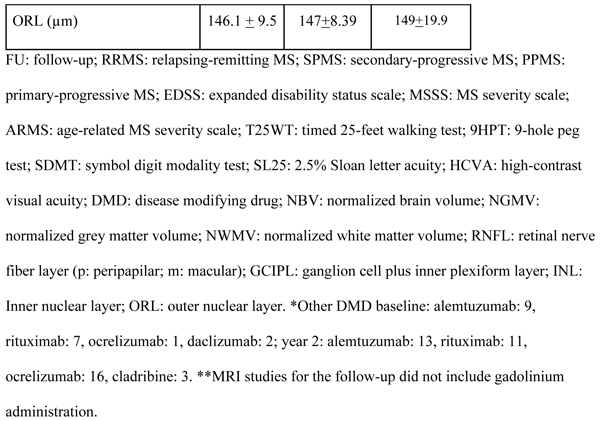
Sys4MS cohort: Clinical and imaging variables of MS patients and healthy controls. Disability scales are shown as the mean ± SD, except for the EDSS which is displayed as the median (range).

|  |  | MS<br>baseline<br>n=328 | MS<br>2-year FU<br>n=278 | HC<br>n=90 |
| --- | --- | --- | --- | --- |
| Age | | 41±10 | 45±9.81 | 36.98 ± 11.47 |
| Female, n (%) |  | 229 (70%) | 194 (70%) | 63 (70%) |
| Age at disease onset (years) | | 31 ± 9 | 31±9 | -- |
| Disease duration (years) | | 10 ± 8 | 12.9±8.16 | -- |
| Subtype | RRMS | 271 | 228 | -- |
|  | SPMS | 28 | 25 |  |
|  | PPMS | 29 | 25 |  |
| EDSS |  | 2.0 (0-8.0) | 2.0 (0-8.0) | -- |
| MSSS | | 3.6 ± 2.2 | 3.25±2.35 | -- |
| ARMS | | 3.9 ± 2.1 | 3.56±2.26 | -- |
| T25WT (sec) | | 6.93 ± 6.6 | 5.67±4.97 | -- |
| 9HPT (sec) | | 21.2 ± 6.5 | 21.9±5.92 | -- |
| SDMT (# symbols) | | 53.8 ± 13.5 | 53.5±13.3 | -- |
| SL25 (# letters) | | 29.1 ± 13.4 | 26.7±13.5 | -- |
| HCVA (LogMAR) | | 0.03 ± 0.36 | -0.11±0.44 | -- |
| DMD | Untreated | 91 | 72 | -- |
| Interferon beta |  | 43 | 19 | -- |

|  |  |  |  |
| --- | --- | --- | --- |
| Glatiramer acetate | 39 | 24 | -- |
| Teriflunomide | 28 | 21 | -- |
| Fingolimod | 38 | 33 | -- |
| Dimethyl-Fumarate | 35 | 37 | -- |
| Natalizumab | 29 | 24 | -- |
| Other high-efficacy DMD* | 19 | 43 | -- |
| Brain MRI |  |  | baseline |
| # Gadolinium lesions | $0.1 \pm 0.5$ | NA** | NA |
| T2 lesion volume (cm<sup>3</sup>) | $8.17 \pm 10.5$ | $9.32 \pm 11$ | NA |
| NBV (cm<sup>3</sup>) | $1,509 \pm 91$ | $1,454 \pm 70.2$ | $1,473 \pm 109$ |
| NGMV (cm<sup>3</sup>) | $792 \pm 65$ | $779 \pm 49.5$ | $751 \pm 63.7$ |
| NWMV (cm<sup>3</sup>) | $716 \pm 68$ | $676 \pm 43.5$ | $721 \pm 111$ |
| OCT (mean of both eyes without previous ON) |  |  | baseline |
| pRNFL (μm) | $100 \pm 12.7$ | $101 \pm 12.1$ | $100 \pm 9.6$ |
| mRNFL (μm) | $39.6 \pm 4.9$ | $39.6 \pm 4.31$ | $41.9 \pm 6.5$ |
| GCIPL (μm) | $65.6 \pm 8.3$ | $65.7 \pm 7.08$ | $68.5 \pm 6$ |
| INL (μm) | $31.5 \pm 2.8$ | $31.5 \pm 2.77$ | $41.1 \pm 8.8$ |
| ORL ($\mu\text{m}$) | $146.1 \pm 9.5$ | $147 \pm 8.39$ | $149 \pm 19.9$ |

FU: follow-up; RRMS: relapsing-remitting MS; SPMS: secondary-progressive MS; PPMS: primary-progressive MS; EDSS: expanded disability status scale; MSSS: MS severity scale; ARMS: age-related MS severity scale; T25WT: timed 25-feet walking test; 9HPT: 9-hole peg test; SDMT: symbol digit modality test; SL25: 2.5% Sloan letter acuity; HCVA: high-contrast visual acuity; DMD: disease modifying drug; NBV: normalized brain volume; NGMV: normalized grey matter volume; NWMV: normalized white matter volume; RNFL: retinal nerve fiber layer (p: peripapilar; m: macular); GCIPL: ganglion cell plus inner plexiform layer; INL: Inner nuclear layer; ORL: outer nuclear layer. \*Other DMD baseline: alemtuzumab: 9, rituximab: 7, ocrelizumab: 1, daclizumab: 2; year 2: alemtuzumab: 13, rituximab: 11, ocrelizumab: 16, cladribine: 3. \*\*MRI studies for the follow-up did not include gadolinium administration.

We conducted a genomic analysis in both MS cases and controls. From the 700,000 SNPs assessed in the DNA array, we imputed 152 SNPs associated with MS (8), along with 17 additional SNPs corresponding to HLA-class II alleles. We calculated the polygenic risk score, namely the MS genetic burden score (26) (MSGB) for all 169 SNPs, together with partial MSGB scores for only the 17 HLA SNPs (MSGB^HLA^), and for the 152 MS associated SNPs excluding the HLA alleles (MSGB^non-HLA^). As expected, the total MSGB score was significantly higher (p=3.4×10^-8^) in patients (4.23) than in HCs (3.2). Similar results were observed in the partial scores, with MSGB^HLA^ of 1.57 in patients and 0.95 in HCs (p=1.6×10^-4^) and MSGB^non-HLA^ of 2.6 in patients and 2.2 in HCs (p=6.8×10^-5^).

Flow cytometry analysis was carried out at baseline in peripheral blood mononuclear cells (PBMCs) from the first 227 patients and 82 HC. Results from the cytometry analysis in this cohort are described in detail elsewhere (24). Briefly, untreated RRMS patients showed significantly higher frequencies of Th17 cells and lower frequencies of B-memory/B-regulatory cells, as well as higher percentages of mature B cells in patients with PMS compared with HCs. Fingolimod treatment induced a decrease in total CD4+ T cells and mature and memory B cells and increases in CD4+, CD8+ T-regulatory and B-regulatory cells (24). Finally, the phosphoproteomic analysis was carried out by conducting ex-vivo assays in PBMCs and quantified using xMAP assays on the first 148 patients at baseline as described before (25, 27), showing higher levels of phosphorylated IKBA, JUN, KSGB1, MK03, RS6, STAT3 and STAT6 in MS patients compared to controls (**Methods, File S1**).

### Multilayer networks in MS

We built networks for each of the five layers (genetics, phosphoproteomics, cytomics, tissue/imaging and clinical variables) using mutual information to define connections between pairs of elements within each layer (**Figure 1**, see **Methods**). For example, in the proteomics layer two proteins are connected to each other with a weight equal to the normalized mutual information between their phosphorylation levels. A threshold was used to determine whether the correlation for a given pair was high enough to define an edge. The threshold works by comparing the real mutual information value of a pair of nodes to a surrogate distribution of mutual information values calculated from random permutations of the data.

The genetic network was considered in two ways: first, at the level of the individual SNPs separately and utilizing previous information from the Gene Regulatory Network Database (28) and mapped to the MS associated SNPs (see **Methods**); and second, grouped together in the three MSGB scores defined above. The proteomic network includes 25 kinases, and the cytomics network 22 immune subpopulations (see **Methods** for the lists of proteins and cell subtypes). The imaging network included the main metrics of lesion load and brain volumes quantified by MRI, and the thickness of the retina layers analyzed by OCT. Finally, the clinical network contains demographic and clinical variables (number of relapses, disability scales and use of DMD) at baseline and after two-year follow-up, which give longitudinal changes in clinical outcomes (see **Methods** for a list of variables).

After the networks for each layer were built, we analyzed the connectivity (density) between layers, this time between features of different layers. A statistical comparison between the connections within and between layers (**Figure 2**) shows a non-negligible degree of network modularity, confirming the underlying multi-layer structure. The features within a layer are, on average, more strongly connected than those between layers. With the exception of genomics, the densities within a single layer were higher than those between layers, supporting the modularity of the multilayer network.

**Figure 2:**
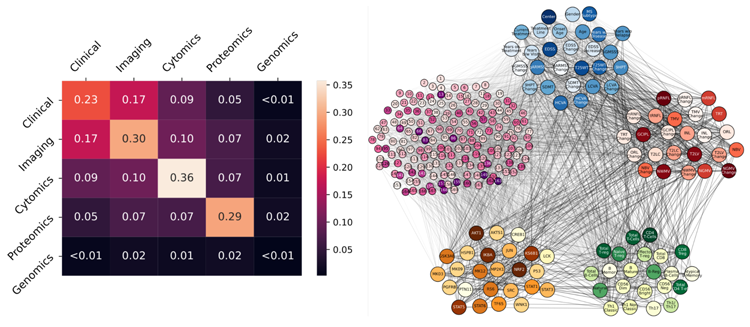
Network densities within and between layers. (*left*) The density for each layer was calculated as the ratio of the sum of the weights of all connections and the number of possible connections. The analysis was made using the 67 subjects with complete data in all 5 layers. (*right*) The network from which the density was calculated. Nodes from all layers were connected together, opposed to the network model with the hierarchy shown before. See high resolution network at https://keithtopher.github.io/combo_networks/<u>#/</u>.

### Dynamic network analysis identifies gene-protein-cell paths associated with phenotype

We next sought to integrate all the layers in paths that reflect the network dynamic interactions, in order to obtain a functional view of the information flow across layers. To that end, we created a single network including all five layers with the same hierarchy described above using linear (Pearson) correlations, which allows us to distinguish between stimulatory or inhibitory edges depending on whether the correlation r value is positive or negative, respectively (**Figure 3a**). We then conducted logic (Boolean) simulations to identify the causal logic backbone of the network (29, 30). Boolean simulations use knowledge of activating and inhibiting relationships between nodes; the exact chemical reactions between genes, proteins, cells and tissue are ignored, giving a qualitative description of the system (30). The nodes of the network are considered to be in one of two states: active (e.g., high phosphorylation levels) or inactive (e.g., low phosphorylation levels). The states of all nodes are updated synchronously at each iteration of the simulation, either remaining in the same activation state as before, or flipping to the opposite state, depending on the activation states of its direct neighbors, and taking into consideration the weights of the corresponding connections (**Figure 3b**, see **Methods**).

**Figure 3:**
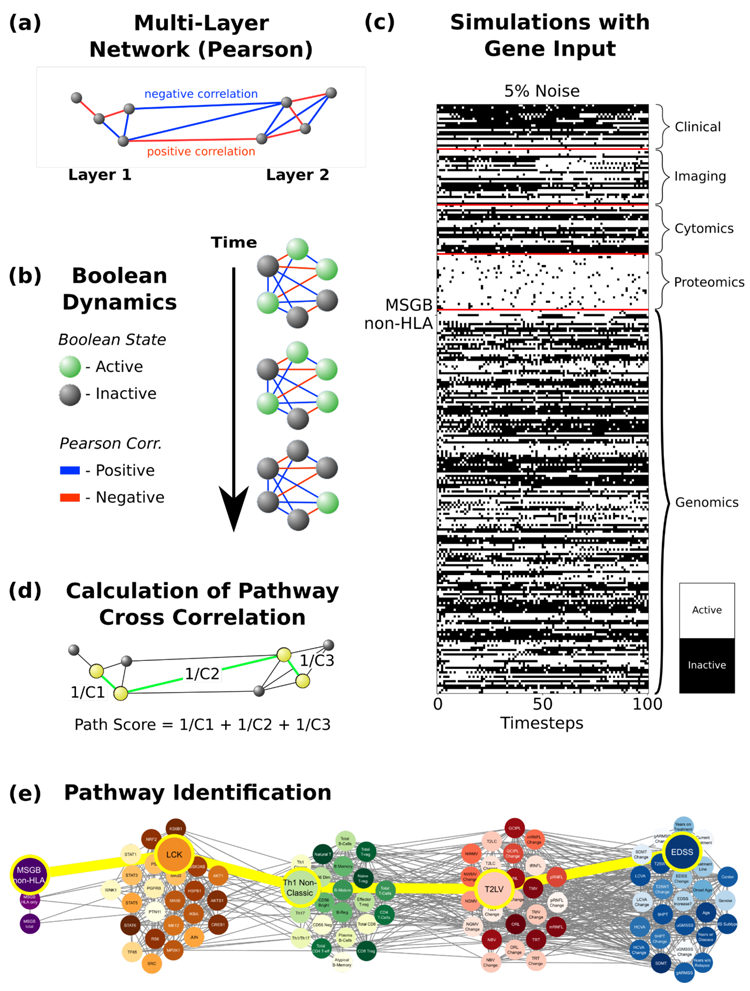
Dynamic network analysis: identification of gene-protein-cell paths. (a) Networks are constructed using all five layers. The nodes are the same as in the networks above (figure 1), but now the edges are defined by the Pearson correlation, where the weights represent the Pearson coefficient, which can be either positive or negative. (b) Boolean dynamics are applied to the networks, where the activation state of the nodes changes based on the total sum of the edge weights of its direct neighbors (considering the signs of the connections). (c) Boolean simulations are run where the various nodes, in the example MSGB non-HLA, are used as the input signal, and the simulation was run with 5% noise (see **Methods** for noise analysis). (d) The cross-correlation coefficient (Cn) is calculated between the signals for each pair of connected nodes. A path score is calculated for all possible paths, defined as the sum of the inverses of the cross-correlation coefficients between all pairs of consecutive nodes constituting a given path. (e) Finally, a path is identified by using a shortest path algorithm which is based on its path score (see **Methods**).

We next wanted to study how perturbations in a given input such as the MSGB score (SNPs could not be used for Boolean simulations because the impossibility of changing between alleles), protein or cell type travel through the network and ultimately affect a given phenotype (output). To that end, we performed Boolean simulations in which the input node was periodically driven from an active to an inactive state and back, and the response of all nodes in the network (**Figure 3c**) was quantified by computing the temporal cross-correlation function between their time-varying state and the dynamic input signal (30). We then identified those paths across the network that are formed by pairs of nodes with the highest temporal cross-correlation between their signals. These paths represent how information flows from a given input to the output (e.g., from MSGB non-HLA to EDSS in the example in **Figure 3e**). They do not necessarily represent physical interactions among nodes (e.g., protein-protein interactions), but rather groups of nodes that co-vary statistically with each other more strongly than the rest of the network.

For each of the 3,350 combinations of inputs and outputs (3 MSGB scores, 25 proteins, and 22 cell types as inputs, and 22 cell types, 25 imaging variables, 20 clinical variables as outputs), we selected the top ten paths with highest joint cross-correlation values between their constituent nodes (see **Methods** and **File S2**). **Figure 4** shows these paths for the three inputs (MSGB, phosphoproteomics and cytomics) and outputs (imaging and clinical) pairs for MS patients.

**Figure 4:**
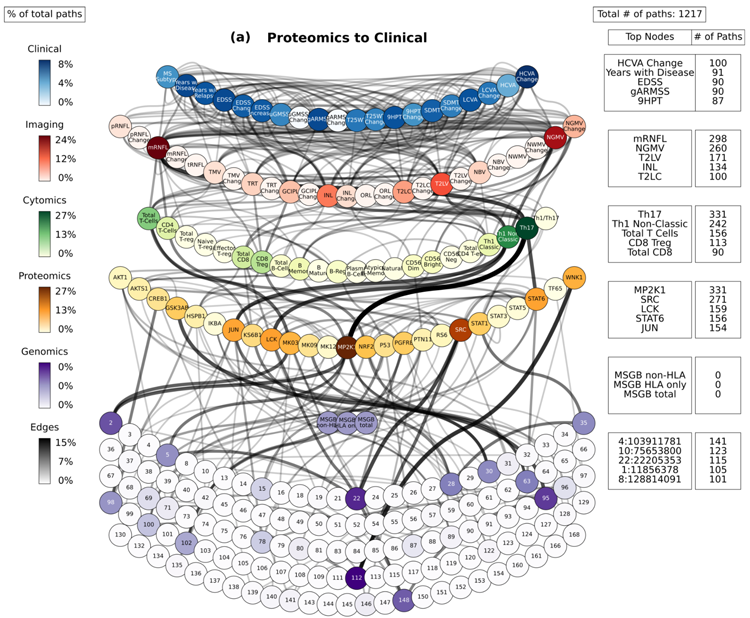
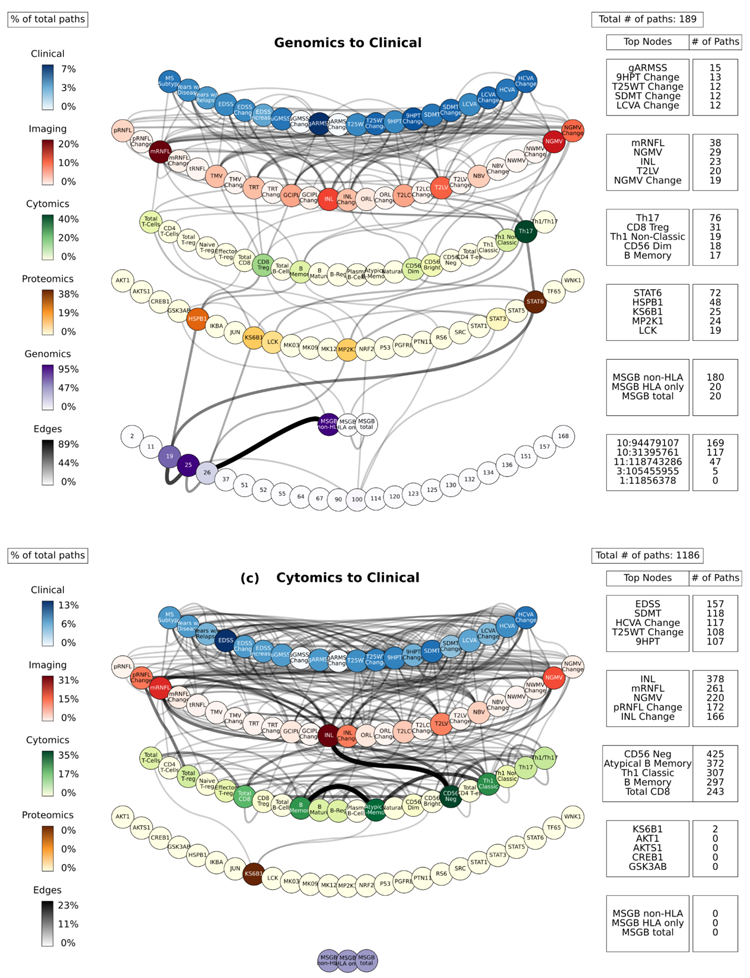
Path analysis in MS patients. Representations of the multi-layer paths identified from the Boolean simulations when the input started at the phosphoproteomics (A), genomics (B) or cytomics (C) layer. The top paths (those that passed the test for negative controls) are shown for each input (gene, protein, or cell)-output (clinical phenotype) pair. The nodes for each layer are color-coded to represent the degree of a given node, i.e., the number of times the node appears in a path, as a percentage of the total number of paths. High resolution paths are available at https://keithtopher.github.io/fivelayer_pathways/.

To assess the specificity of the Boolean simulations, the network was permuted to identify negative control paths. The edges were randomly swapped while preserving the original degree distribution of the network. The simulations were run with these permuted networks (100 total), and paths were identified. These paths were compared to those identified in the original networks. We counted how many times a given path appeared in the permuted networks. Focus was placed on those pathways that were present in less than 1% of the permuted pathways. Out of 32,302 total paths identified from MS patients, there were 8,488 that appeared 0 times out of 100 in the permuted paths. The method for network permutation and path identification is illustrated in **Methods, Figure 10**, and results are shown as **Files S3a and S3b**.

Additionally, confidence intervals were calculated for each of the paths. The paths were identified from each of the 100 Boolean simulations individually (instead of using the mean of the cross-correlation values as before). These 100 simulations provide a distribution of path scores, giving the variance of the original path score. The path scores along with their confidence intervals are given in **File S4**.

### Path analysis

For the path analysis we use the following notation: NODE 1 > NODE 2 > NODE 3, (In the case there are multiple nodes on the same layer along similar paths they appear as NODE 1 > NODE 2 - NODE 3 > NODE 4, where NODES 2 and 3 could be two proteins for example.) and the information flows from left to right, starting with the perturbation in the gene, protein, or cell respectively. MS cases show that the paths more commonly found from the Boolean simulations (darker color represents more connections) were: (1) MP2K1 > Th17 > mRNFL > ARMSS (when the input is applied to the started in phosphoproteomics layer, **Figure 4a**); (2) SNP25 (SNP10:94479107) > MSGB non-HLA > STAT6 > Th17 > mRNFL> ARMSS (when the input is applied to the genomics layer, **Figure 4b**); (3) CD56 Neg > INL - mRNFL > EDSS - ARMSS (when the input is applied to the cytomics layer, **Figure 4c**).

Perturbations in the protein layer (representing changes in the signaling cascades among cells) were linked with the severity of MS, this time with both the EDSS and ARMSS along with the HCVA, T25WT and the disease duration (**Figure 4a**):

- MK03 > Total T Cells > mRNFL > T25WT
- MP2K1 - STAT6 > Th17 > mRNFL > T25WT - ARMSS
- MP2K1 - STAT6 > Th17 > INL > EDSS Change
- MP2K1 > CD8 Treg > GCIPL > EDSS Change
- LCK - JUN > Th1 non-Classic > NGMV - mRNFL - T2LV
- NGMV > Years since Relapse - 9HPT Change - HCVA Change mRNFL > T25WT - Years with Disease - SDMT Change T2LV > EDSS - ARMSS - T25WT
- SNP10:75653800 - SNP4:103911781 - SNP1:85729820 > SRC - STAT6 - AKTS1 - NRF2

Perturbations of the gene network (the MSGB, reflecting genetic variability contributing to the risk of developing MS) were linked with changes in the clinical outcomes (ARMSS, T25WT, 9HPT, HCVA, LCVA, and the EDSS) (**Figure 4b**). Concerning the imaging layer, we found paths to the mRNFL (macular retinal nerve fiber layer) and NGMV (normalized gray matter volume). Perturbing the MSGB non-HLA was the source for the most paths at this level:

1. MSGB non-HLA > SNP10:94479107 > SNP11:118743286 > KS6B1 - MP2K1; and 2) MSGB non-HLA > SNP10:94479107 > SNP10:31395761 > HSPB1 - STAT6. Then, these two paths were connected to the phenotype as follows:

- HSPB1 > B Memory > NBV > MSSS - T25WT
- HSPB1- MP2K1 > CD8 Treg - B Memory > GCIPL > ARMSS - EDSS Change
- STAT6 > Th17 > mRNFL - INL > ARMSS
- STAT6 > Th17 > NGMV Change > Years with Disease
- MP2K1 > Th17 - CD8 Treg > mRNFL - INL > ARMSS - EDSS Change - 9HPT Change
- KS6B1 - LCK > Total T Cells - Th1 Non Classic > NGMV - T2LV > LCVA Change - MSSS - Years since Relapse

Perturbations at the cellular level (representing changes of immune cell subtypes frequency and activation) were connected again with changes in the EDSS as well as with the HCVA, SDMT, 9HPT, and T25WT (**figure 4c**). The paths with cells as the sources were:

- CD56 Neg > INL - mRNFL > EDSS - T25WT
- Atypical B Memory - B Memory - Th1 Classic > mRNFL - T2LV > EDSS - T25WT
- Total CD8 > NGMV - T2LV > EDSS - 9HPT - SDMT

### Paths predicting MS phenotype from single-cell data

In order to assess some of the paths identified in the study at the single-cell level, we conducted a cytometry analysis to assess levels of total and phosphorylated proteins in immune cell subtypes at the single-cell level and relate them to the clinical phenotype through linear regression models and path analysis. We analyzed the levels of the three phosphoproteins for which phospho-cytometry assays were available and that showed an adequate signal to noise ratio, namely GSK3AB, HSBP1 and RS6 (assays were not validated for the other proteins). We also analyzed the immune cell subpopulations most commonly present in such paths (CD4+, Treg, CD8+, B mature, B memory, Breg and Plasma cells). This approach allowed to assess experimentally the paths between an individual phosphoprotein in the selected immune cell subtype. The phosphorylation levels in immune subpopulations were assessed in a representative subgroup of 40 MS patients and 20 HCs from the Sys4MS cohort from which frozen PBMCs were available from the baseline visit (**Figure 7**).

First, we found significant linear regression models for each of the three kinases predicting the phenotype (**Figure 5**) (see **Files S5a, S5b, and S5c** for R^2^ and p-values). In the case of GSK3AB, we found significant regression models explaining disease duration, walking speed, retina, and grey matter atrophy. For HSPB1, significant regression models were found for global disability scales such as the EDSS as well as domain specific disability scales (motor, vision, cognition), disease duration and change in grey and white matter volume. Finally, for RS6 the significant regression models also explained changes in global and motor disability (GMSSS and 9HPT) as well as retina and brain atrophy.

**Figure 5.**
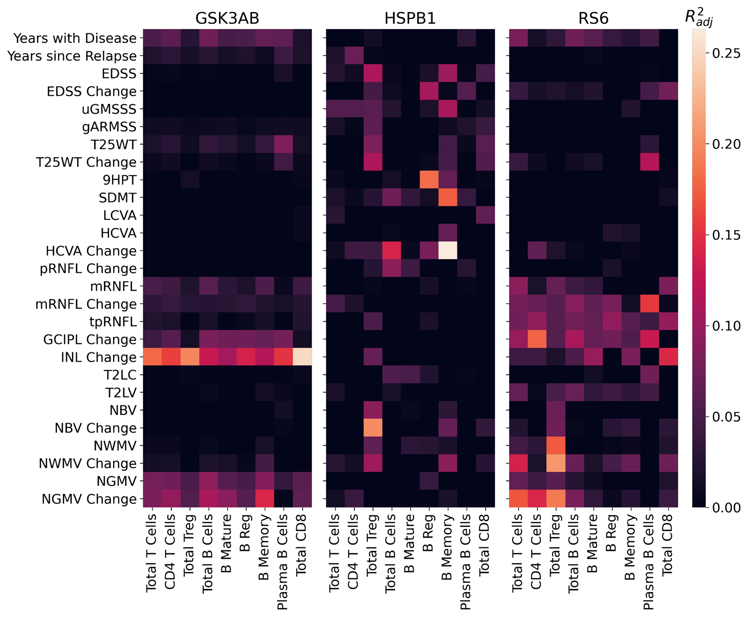
Linear regression models between phosphoproteins, cell subtypes and clinical phenotype. Linear regression analysis relating the percentage of immune cell subtypes expressing phosphorylated GSK3AB, HSBP1 or RS6 with the phenotype. The heatmap shows the adjusted R^2^ of the significant models. EDSS: Expanded Disability Status Scale; GMSSS: Global Multiple Sclerosis Severity Score; T25WT: timed 25 feet walking test; 9HPT: nine-hole peg test; LCVA: low contrast (2.5%) visual acuity; HCVA: high contrast visual acuity; RNFL: retinal nerve fibber layer (m: macular; tp: temporal peripapillary); INL: inner nuclear layer; T2LV: T2 lesion volume; ORL: outer retinal layer; NBV: normalized brain volume; NWMV: normalized white matter volume; NGMV: normalized grey matter volume.

We then applied the single-cell data to our multilayer network and paths shown in **Figure 4**. The network was made using the significant values from the linear regressions to relate phosphoprotein-cell layer to the phenotype. With each protein (GSK3AB, HSPB1, RS6), wherever there was a significant value between a cell and phenotype, an edge was placed between the protein and cell, and another edge between the cell and the phenotype. For example, there is a significant model between the percentage of B Memory cells expressing GSK3AB and the INL change, so the two edges GSK3AB > B Memory and B Memory > INL Change are added. Edges between the imaging and clinical layers are formed indirectly, where two nodes are connected if they had at least one significant regression model with the same cell type. For example, since there are significant models between Total Treg and NBV, as well as between Total Treg and EDSS, the edge NBV > EDSS is added. Next, edges between the cellular and clinical layers are removed. Finally, only the edges that are also found in the top paths from the five-layer network shown in **Figure 4** are kept. The top paths beginning with GSK3AB, HSPB1, and RS6 are listed below, and a visualization of the paths is shown in **Figure 6**.

- GSK3AB > Total B Cells - Total T Cells - Total Treg > INL Change > Years with Disease - EDSS Change
- GSK3AB > Total B Cells - Total T Cells - Total Treg > NGMV Change > Years with Disease
- GSK3AB > Total T Cells - Total Treg > NWMV Change > Years with Disease - EDSS - GMSSS - T25WT
- GSK3AB > Total B Cells - Total T Cells > tRNFL > Years with Disease - EDSS Change
- HSPB1 > Total B Cells - Total Treg > INL Change > Years with Disease - EDSS Change
- HSPB1 > Total B Cells - Total Treg > NGMV Change > Years with Disease
- RS6 > Total B Cells - Total T Cells - Total Treg > INL Change > Years with Disease - EDSS Change
- RS6 > Total B Cells - Total T Cells - Total Treg > NGMV Change > Years with Disease
- RS6 > Total T Cells - Total Treg > NWMV Change > Years with Disease - EDSS - GMSSS - T25WT
- RS6 > Total B Cells - Total T Cells > tRNFL > Years with Disease - EDSS Change

**Figure 6:**
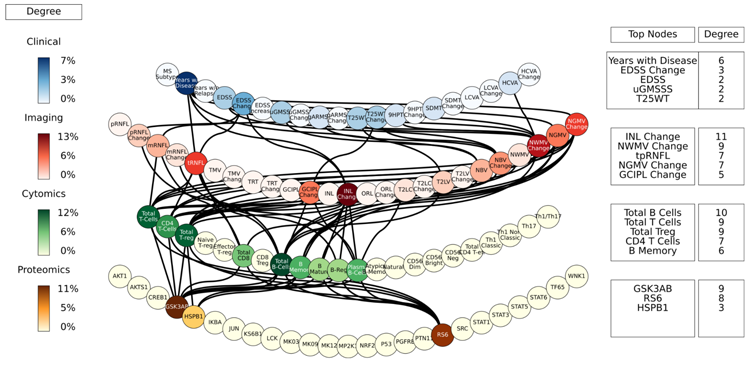
Multilayer paths from single cell cytometry assays. Each of the edges was defined using the linear regression analysis of the flow cytometry data. An edge is considered if it was part of a significant regression model and also appeared as part of a path in the original five-layer network constructed from MS patient data (from Figure 4). The edges are weightless, and only show if that particular edge in any of the original paths was present. https://keithtopher.github.io/fivelayer_pathways/.

**Figure 7.**
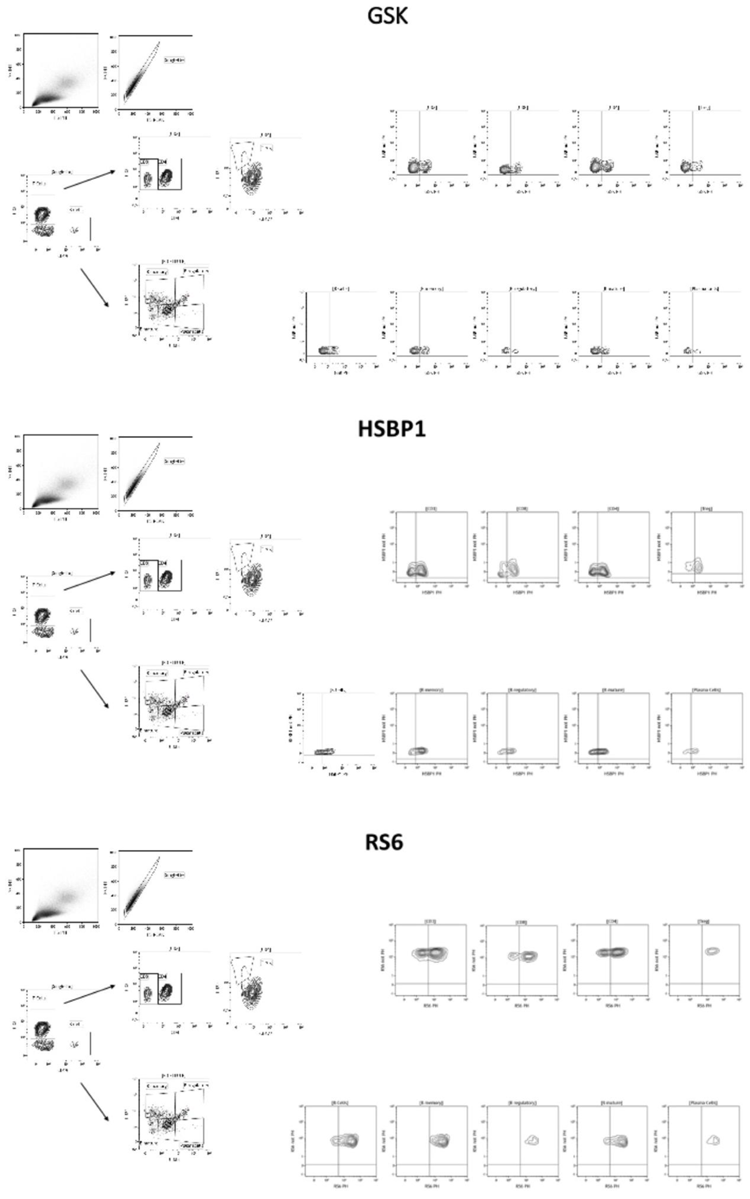
Cytometry plots for the expression of phosphoGSK3AB, phosphoHSBP1 and phosphor RS6 in immune cell subpopulations. The gating strategy for phospho-flow cytometry analysis. Examples of phospho-GSK3AB, phospho-HSBP1, and phospho-RS6 staining in the immune cell subpopulations for MS patients are presented.

## Discussion

Network approaches have been very fruitful in the past at shedding light on the molecular complexity of diseases, beyond the traditional single gene and single pathway perspectives. In the traditional network paradigm, molecular components are connected according to their biological interactions, and the structure and dynamics of such interaction networks can reveal disease modules and nonlinear pathways (32). Recently, these approaches have been extended to include multiple biological layers, such as diverse tissues with distinct protein-protein interaction networks (33), and different biological processes (membrane potential dynamics and signaling) within insulin-secreting cells (34). Attempts have been made to construct multilayer networks for complex diseases, an approach successfully exploited in cancer research (35–40). In this study we have applied a multilayer network analysis to integrate omics, imaging, and clinical information from patients with a complex autoimmune disease such as MS.

Our multilayer network analysis allowed us to assess the relationship between different biological scales in the disease and to identify paths linking the five layers (genomics, proteomics, cytomics, imaging and clinical) based on statistical associations. The most relevant multiscale paths from our study are:

1. MP2K1 > Th17 > mRNFL > ARMSS;
2. SNP25 > MSGB non-HLA > STAT6 > Th17 > mRNFL> ARMSS;
3. CD56 neg > INL - mRNFL > EDSS - T25WT.

The interaction of several phosphoproteins-cell paths and the phenotype were validated by flow cytometry studies, which were based on single cell analysis. A multi-layer network analysis is thus able to identify a differential activation of the immune system’s multiple scales in MS patients that drives the phenotype.

It is of course possible that there were changes on the protein and genetics level, but they were not acting as mediators between the changes in the cell counts and the phenotype seen in this case. These could be considered sub-level systems that may cause the changes in the higher levels when concerning the phenotype.

The results from the multilevel network analysis with the omics data and phenotype data highlight the importance of considering MS as a multiscale disease, where the layers connect with varying strengths and information is filtered or strengthened across the layers (34, 39). Previous studies attempted to directly link the genomic layer with the phenotypes in many complex diseases, including MS. However, genotypes or the polygenic risk scores alone have a limited ability to predict either the cell variability or the phenotype (31, 41). Other genetic information such as DNA sequencing, epigenetics and RNA expression, or more global approaches is likely needed for a more thorough analysis in multiscale complex diseases.

The kinases studied are part of pathways previously described as associated with MS (reviewed in (9)). MP2K1 was the kinase showing the strongest association with the presence of MS in our previous study (25) and is a master regulator of the immune response. We and others have previously described increased GSK3AB expression or phosphorylation levels in patients with MS (25, 42, 43). GSK3 plays key roles in Th1 cell activation as well as in microglia modulation, in addition to its effects on neuronal survival and functioning (42). HSBP1 (also known as HSP27) is a stress protein that in addition to its chaperone activity, is critical for apoptosis signaling pathways within the mitochondria, inhibiting the Apaf complex (44). Indeed, HSBP1 has been found to be increased during MS relapses (45). RS6 is a MAPKinase that is modulated by extracellular signal-regulated kinase (ERK) and activates serum glucocorticoid kinase 3 (SGK3), nuclear factor kappa-light-chain-enhancer of activated B cells (NfKB), mammalian target of rapamycin (mTOR) and other pathways modulating cell growth and differentiation. Inhibition of ERK and RS6 in models of MS reduces proliferative response, phagocytic properties, and synthesis of proinflammatory mediators induced by the addition of inflammatory stimuli to microglia (46). Regarding the immune cell subtypes highlighted in our analysis, our previous analyses of the Sys4MS dataset support the results of the current network analysis that confirms the prominent role of B cells in MS (24). Such results agree with our previous analysis of phosphoproteins and immune cell subtypes in another dataset of MS patients showing the preferential involvement of B cells (25). Many pieces of evidence have confirmed a remarkable role for B cells in MS (47), probably driven by the latent infection of the Epstein-Barr virus that produces immune response dysregulation or molecular mimicry with CNS proteins like GlialCAM (16, 48). In addition, CD8 cells are the most abundant cell type in the brain infiltrates (11). Finally, a recent study in twins discordant for MS is also providing new endorsement of the role of helper CD4 cells (12).

The data provided by the Sys4MS cohort was rich in the wide range of scales it covered. However, several limitations were encountered with both the data and analysis. Although the sample size of the cohort was enough to identify significant correlations, the sample sizes were smaller for some specific omics (proteomics and cytomics), although bigger than n-of-1 studies commonly used for deep phenotyping (39). The limited sample size may have affected both the networks constructed as well as the statistical tests conducted with the paths or for the analysis stratifying by each of the therapies. Furthermore, the omics dataset collected were cross-sectional, whereas the imaging and clinical data were longitudinal. Longitudinal data from all five layers and deep phenotyping would greatly benefit future studies. Another concern is the validation of the paths because deep phenotyped MS cohorts are not available. A wealth of MS patient data from other studies is available with genomics, imaging, and clinical phenotype (through the IMSGC and MultipleMS consortia). However, proteomics, cytomics or other types of omics data is usually lacking, which limits conducting validation in independent datasets. Further limitations relate to the omics experiments themselves. Both the protein and cell analyses were conducted using PBMCs, rather than in immune cells from the central nervous system. Also, the protein analysis was not performed at the single cell level but in bulk PBMCs in the overall cohort. Therefore, single-cell dynamics were not captured in the first experiment. However, flow cytometry analysis performed for the validation study provided single-cell information which supports the validity of the findings. Additionally, limitations were also partially balanced by using a hypothesis-driven design that included kinases and cells previously described as differentially activated in MS.

In summary, this study examined the functional connections among various scales of biological data of a complex disease with a complex genetic basis, namely MS. Our multilayer networks support that information flow across scales. This highlights the importance of the molecular and cellular scales when considering explaining the phenotypes of complex diseases. Indeed, these paths could be the target of a future treatment of personalized medicine in MS. This could also be transferable to other autoimmune disorders, commonly sharing disease underlying mechanisms.

## Methods

### Ethical Statement

The Sys4MS project was approved by the Institutional Review Boards at each participating institution: Hospital Clinic of the University of Barcelona, IRCCS Ospedale Policlinico San Martino IRCCS, Oslo University Hospital, and Charité - Universitätsmedizin Berlin University. The Barcelona MS cohort study was approved by The Ethic Committee of Clinical Research, Hospital Clinic Barcelona. Patients were invited to participate by their neurologists, and they provided signed informed consent prior to their enrollment in the study. De-identified data were collected in a REDCap database at the Barcelona center.

### Patients

#### Sys4MS cohort

We recruited a cohort of 328 consecutive MS patients according to 2010 McDonald criteria (49) and 90 healthy controls (HC) at the four academic centers: Hospital Clinic, University of Barcelona, Spain (n=93); Ospedale Policlinico San Martino, Genova, Italy (n=110); Charité - Universitätsmedizin Berlin, Germany (n=94); and the Department of Neurology, Oslo University Hospital, Norway (n=121) as described before (24). We collected clinical information (demographics, relapses, disability scales, and use of disease-modifying drugs), and imaging data (brain MRI and OCT), and obtained blood samples at the same visit. Patients were required to be stable in their DMD use over the preceding six months. Patients were followed for two years, and the same clinical, disability scales, and imaging data (brain MRI and OCT) were collected at the 2-year follow-up visit.

#### Clinical Variables

Each patient was assessed on the following disability scales at baseline and follow-up: the Expanded Disability Status Scale (EDSS); timed 25 feet walking test (T25WT), nine-hole peg test (9HPT), the Symbol Digit Modality Test (SDMT), 2.5% low contrast visual acuity (SL25), and high contrast vision (HCVA, using EDTRS charts and a logMar transformation). We calculated the MS Severity Score (MSSS) and the age-related MS Severity Score (ARMSS).

The ARMSS was used for dividing the cohort based on disease severity using the tertile distribution (first tertile were mild MS, the second tertile was excluded and the third tertile were defined as severe MS). Change in the disability scales and 2-year follow-up visit was calculated as the difference (delta) between the two visits. EDSS changes were confirmed in a clinical visit 6 months before the study follow-up visit. At each visit, we collected the information regarding the patients’ DMD use, including low-efficacy therapy: interferon-beta, glatiramer acetate, and teriflunomide; or mid to high-efficacy therapy: fingolimod, dimethyl-fumarate, natalizumab, or other monoclonal antibodies (alemtuzumab, rituximab, daclizumab, and ocrelizumab).

#### Imaging

MRI studies were performed on a 3-Tesla scanner at each center using a standard operating procedure (SOP) to optimize the volumetric analysis. We used the 3-dimensional (3D) isotropic T1-weighted magnetization-prepared rapid gradient echo (T1-MPRAGE) (resolution: 1 x 1 x 1 mm^3^), and 3D T2-fluid-attenuated inversion recovery (T2-FLAIR) images with the same resolution to quantify changes in brain volume. Presence of contrast-enhancing lesions, T2 lesion volume, new or enlarging T2 lesions, and volumetric analysis were done at the Berlin center as previously described (50, 51).

Retinal OCT scans were performed using the Spectralis device in three centers and the Nidek device at Oslo center. A single grader at the reading center in Berlin performed intra-retinal layer segmentation using Orion software (Voxeleron Inc, Berkeley, US) to quantify the macular ganglion cell plus inner plexiform layer (GCIPL) and the macular inner nuclear layer thicknesses (μm) in the 6 mm ring area as previously described (52).

#### Brain Magnetic Resonance Imaging

All images were acquired from 4 centers with distinct 3-tesla systems after standardizing the acquisition protocols and validating dummy scans by the MRI reading center in Berlin. From Center 1 (Barcelona), a three-dimensional (3D) magnetization prepared rapid gradient echo (MPRAGE) sequence, including the upper cervical cord (0.86 x 0.86 x 0.86 mm resolution, repetition time (TR)=1970 ms, echo time (TE)=2.41 ms), an axial T1-weighted post-gadolinium contrast agent sequence (0.31 x 0.31 x 3 mm resolution, TR=390 ms, TE=2.65 ms), and a 3D fluid-attenuated inversion recovery (FLAIR) sequence, including the upper cervical cord (1 x 1 x 1 mm resolution, TR=5000 ms, TE=393 ms) were acquired longitudinally (2 visits) from 60 MS patients using a Tim Trio MRI (Siemens Medical Systems, Erlangen, Germany). From Center 2 (Oslo), a 3D sagittal brain volume (BRAVO) sequence for pre-and post-gadolinium contrast agent administration, including the upper cervical cord (1 x 1 x 1 mm resolution, TR=8.16 ms, TE=3.18 ms), and a 3D FLAIR sequence, including the upper cervical cord (1 x 1 x 1.2 mm resolution, TR=8000 ms, TE=127.254 ms) were acquired longitudinally (2 visits) from 97 MS patients using a Discovery MR750 MRI (GE Medical Systems,). From Center 3 (Berlin), a 3D sagittal MPRAGE sequence, including the upper cervical cord (1 x 1 x 1 mm resolution, TR=1900 ms, TE=3.03 ms), and a 3D FLAIR sequence, including the upper cervical cord (1 x 1 x 1 mm resolution, TR=6000 ms, TE=388 ms) were acquired longitudinally (2 visits) from 87 MS patients using a Tim Trio MRI (Siemens Medical Systems, Erlangen, Germany). From Center 4 (Genova), a sagittal fast-spoiled gradient-echo (FSPGR) sequence, including the upper cervical cord (1 x 1 x 1 mm resolution, TR=7.312 ms, TE=2.996 ms), a 3D turbo field echo (TFE) sequence for post-gadolinium contrast agent administration (1 x 1 x 1 mm resolution, TR=8.67 ms, TE=3.997 ms), and a 3D FLAIR sequence, including the upper cervical cord (1 x 1 x 1 mm resolution, TR=6000 ms, TE=122.162 ms) were acquired longitudinally (2 visits) from 88 MS patients using a Signa HDxt MRI (GE Medical Systems) and Ingenia MRI (Philips Medical Systems).

#### MRI Post-processing

Analysis for all scans were conducted at the MRI reading center in Berlin. Preprocessing included registration to MNI-152 standard space (fslreorient2std), white and grey matter brain masking (Computational Anatomy Toolbox 12 Toolbox for MATLAB SPM12, http://www.neuro.uni-jena.de/cat/), N4-bias field correction (Advanced Normalization Tools, http://stnava.github.io/ANTs/) and linear, rigid body registration of T2-weighted (FLAIR) images to T1-weighted (MPRAGE, BRAVO, and FSPGR) images (FSL FLIRT, https://fsl.fmrib.ox.ac.uk/fsl/fslwiki/FLIRT/UserGuide). Each second session for each patient T1-weighted image and FLAIR image was co-registered to the individual first session using the transformation matrices saved from the first session transformation from native space images to MNI-152 standard space using FSL FLIRT. Post-contrast agent T1-weighted images were also co-registered to MNI-152 standard space and longitudinally when available.

#### Brain Lesion Segmentation

T2-hyperintense lesion segmentation was performed manually on co-registered T1-weighted images and T2-weighted FLAIR images by two experienced MRI technicians from the Berlin center. Lesions were segmented and saved as binary masks using ITK-SNAP (www.itksnap.org). First session lesion masks were subsequently overlayed onto second session co-registered T1-weighted and FLAIR images for editing, to include any T2-hyperintense lesion changes (i.e., new lesions, enlarging lesions, or decreasing lesions) in the follow-up scans. Any discrepancies in co-registrations that were visible between sessions were corrected manually using the ITK-SNAP automated registration tool prior to follow-up lesion mask edits. Binary gadolinium enhancing lesion masks were created manually using the same tools on the post-gadolinium T1-weighted MR images by the same two technicians. Lesion counts and volumes were extracted from lesion masks using FSL maths (https://fsl.fmrib.ox.ac.uk/fsl/fslwiki/Cluster).

#### MRI Analysis

T2-hyperintense lesion masks were used to fill longitudinally co-registered T1-weighted (not post-gadolinium scans) images using FSL lesion filling (https://fsl.fmrib.ox.ac.uk/fsl/fslwiki/lesion_filling) with white matter masks created from the Computational Anatomy Toolbox for SPM12 (CAT12, http://www.neuro.uni-jena.de/cat/). Lesion filled T1-weighted images were then used for whole brain white and grey matter volume extraction, including the follow-up session percent brain volume change (PBVC) using FSL SIENAX/SIENA (https://fsl.fmrib.ox.ac.uk/fsl/fslwiki/SIENA). The same T1-weighted lesion-filled images were used for whole thalamus volume (sum of left and right thalamic volumes) calculation using FSL FIRST (https://fsl.fmrib.ox.ac.uk/fsl/fslwiki/FIRST). All volumes are reported in milliliters.

#### Optical Coherence Tomography

Retinal OCT scans were performed using the Spectralis device in three centers and the Nidek device at Oslo center. OCTs were collected in eye-tracking mode by trained technicians under standard ambient light conditions (lighting level of 80–100 foot-candles) and without pupillary dilatation. Correction for spherical refractive errors was adjusted prior to each measurement, and the technicians performing OCT scans were aware of the patient’s clinical history. The peripapillary Retinal Nerve Fiber Layer thickness (pRNFL, μm) was measured with a 12-degree diameter ring scan automatically centered on the optic nerve head (100 ART, 1,536 A-scans per B scan). The macular scan protocol involved a 20 x 20-degree horizontal raster scan centered on the fovea, including 25 B scans (ART ≥9, 512 A-scans per B scan). A single grader at the reading center in Berlin performed intra-retinal layer segmentation using Orion software (Voxeleron Inc, Berkeley, US) to quantify the macular ganglion cell plus inner plexiform layer (GCIPL) and the macular inner nuclear layer thicknesses (μm) in the 6 mm ring area as previously described (52). All OCT scans fulfilled OSCAR-IB criteria and scans with an insufficient signal to noise ratio, or when the retinal thickness algorithm failed were repeated, or the data was ultimately excluded.

#### Flow cytometry

The original cytometry data was obtained on fresh peripheral blood mononuclear cells (PBMCs) using 17 antibodies that covered 22 cell subpopulations of T, B and NK cells as described in detail elsewhere (24). The following cell populations were studied: T cells: CD3+, CD3+CD4+, CD3+CD8+; B cells: CD19+; and NK cells: CD3-CD14-CD56+, as well as the specific subpopulations: Effector cells: Th1 classic: CD3+CD4+CxCR3+CCR6-CD161-; Th17: CD3+CD4+CxCR3+CCR6-CD161+CCR4+; Th1/17: CD3+CD4+CCR6-CD161+CxCR3highCCR4low; Regulatory T cells: CD3+CD4+: Treg CD25+CD127-, T naive CD45RA+CD25low; CD3+CD8+: T reg CD28-and T naive CD28-CD45RA+; B cells: B memory: CD19+CD14-CD24+CD38-; B mature: CD19+CD14-CD24+CD38low; B regulatory: CD19+CD24highCD38high and NK cells: Effector: CD3-CD14-CD56dim: Regulatory: CD3-CDCD56bright (reg). For validation assays, PBMC in triplicate tubes were stained with BV510-conjugated anti-CD3 (Clone OKT3, Catalog # 317332, BioLegend)), APC Cy7-conjugated anti-CD4 (Clone SK3, catalog #344616, BioLegend), BV421-conjugated anti-CD25 (Clone BC96, catalog # 302630, BioLegend), AF700-conjugated anti-CD127 (Clone A019D5, catalog # 351344, BioLegend), PE Cy7-conjugated anti-CD19 (Clone HIB19, catalog # 302215, BioLegend), PE-conjugated anti-CD24 (Clone ML5, catalog # 311105, BioLegend) and PE/Dazzle594-conjugated anti-CD38 (Clone HB-7, catalog # 356630, BioLegend) antibodies in solution for 30 min at 4° C and washed twice with PBS. The cells were then fixed and permeabilized with Cytofix/Citoperm (BD Bioscience), according to the manufacturer’s instructions. For intra-cellular staining, the cells were blocked with 5% normal goat serum for 20 min on ice to prevent non-specific binding of the antibodies, and stained for total and relevant phosphoproteins with the following antibodies in one of the three tubes: Tube 1: mouse monoclonal anti-human RPS6 (Clone 522731, catalog # MAB5436, R&D Systems) and rabbit polyclonal anti-human Phospho-RPS6 (Catalog # AF3918, R&D Systems); Tube 2: rat monoclonal anti-human GSK-3B(Clone 272536, catalog # MAB2506, R&D Systems) and rabbit polyclonal anti-human Phospho-GSK-3BCatalog # AF1590, R&D Systems); and Tube 3: mouse monoclonal anti-human HSP27 (Clone G31, catalog # 2402; Cell Signaling Technology) and rabbit polyclonal anti-human Phospho-HSP27 (Catalog #AF2314, R&D Systems) antibodies. All primary antibodies were used at a concentration of 5 7g per 1 x 106 cells. The cells were then washed twice and incubated on ice for 15-20 min with the appropriate fluorescent-conjugated secondary antibodies, Alexa Fluor 488-conjugated goat anti-rabbit IgG (Catalog # A-11070, Invitrogen; 1:100 dilution), APC-conjugated goat anti-mouse IgG (Catalog # 405308, BioLegend; 1:100 dilution), or APC-conjugated goat anti-rat IgG (Catalog # 405407, BioLegend; 1:100 dilution), in 5% normal goat serum. The cells were then washed twice, resuspended in assay buffer, and analyzed on a Beckman Coulter Navios flow cytometer. Analysis was performed using Kaluza software. Phosphorylation levels were defined in terms of mean fluorescence intensity (MFI) of phosphorylated protein over MFI of total protein. A representative cytometry plot for each of the three phosphoproteins is shown in Figure 7. Gating strategy and representative cytometry plots for showing the cell sorting and signal intensity for phospho-GSK3Ab, phospho-HSBP1 and phospho-SR6 assays.

#### Genotyping

Genotyping of the samples was performed by FIMM Genomics (University of Helsinki, Finland) on the Illumina HumanOmniExpress-24 v1.2 array (713,599 genotypes from 396 samples). SNPs imputation was conducted against the 1000-genomes reference (quality of imputation r^2^ > 0.5; 6,817,000 genotypes for 396 samples), which allowed to extract MS-associated SNPs (152 out of 200 known MS-associated SNPs available and 17 out of 31 known MS-associated HLA alleles available (HLA*IMP program)) as described elsewhere (53). The MS Genetic Burden Score (MSGB) for the HLA and non-HLA alleles and their combination was calculated as described previously (26). Briefly, the MSGB is computed based on a weighted scoring algorithm using one SNP per MS associated genomic region as found by trend-test association (meta-) analysis. This statistic is an extension of the log additive model, termed “Clinical Genetic Score”, with weights given to each SNP based on its effect size as reported in the literature. The MSGB is obtained by summing the number of independently associated MS risk alleles weighted by their beta coefficients, obtained from a large GWAS meta-analysis, at 177 (of 200) non-MHC (major histocompatibility complex) loci and 18 (of 32) MHC variants, which includes the HLA-DRB1*15:01-tagging single-nucleotide polymorphism (SNP) rs3135388.

#### Expanded genetic network including regulatory network information

The SNPs were mapped with their nearest gene by the IMSVISUAL consortium (59), and a network was constructed using data from the Gene Regulatory Network Database (GRNdb) (60, 61). The database provides networks of transcription factors (TFs) from various cell types in the human body. The gene regulatory network (GRN) within PBMCs was used containing 12,878 genes, of which we only considered the subset of genes that were mapped to the SNPs from our study. Taking a subset in this way causes some of the regulatory information to be lost, such as two genes that are regulated by the same TF. There is still a relationship between two such genes, although indirect. To include this information in the network of MS genes, an edge was added between two genes that share a transcription factor.

Once the GRN of MS genes was obtained, each gene was then replaced with its corresponding SNP. This is not a one-to-one mapping, as there are some SNPs that are mapped 13 to the same gene. In this case, edges are placed among all SNPs that share a gene. This allows the GRN to be compared with the other layers in the combined network. Finally, only edges that appear in the original network of SNPs connected with Pearson correlation are kept, and their weights are used in the GRN. Details of these networks can be found in https://keithtopher.github.io/networks/<u>#/</u>.

#### XMAP Phosphoproteomics

Phosphoprotein levels were quantified using xMAP assays performed blindly at ProtAtOnce (Athens, Greece) as described previously (25, 27). We analyzed a set of kinases associated with MS (9) which provides an adequate signal to noise ratio and test-retest reproducibility: AKT1, AKTS1, CREB1, GSK3AB, HSPB1, IKBA, JUN, KS6B1, LCK, MK12, MK03/01, MK09, MP2K1, NRF2, P53, PGFRB, PTN11, RS6, SRC, STAT1, STAT3, STAT5, STAT6, TF65, WNK1. Phosphoprotein data was normalized after the measurements were taken as described elsewhere (27).

#### Data Processing

The omics and clinical datasets were ultimately used to build the multilayer network, where each dataset represents a layer in the network. The data were examined to handle missing values, identify which patients have data from which layers, as well as divided into groups based on gender, disease severity, medication, etc. No imputation was used in this study. Patients were divided into mild and severe groups according to the tertiles of their age-related multiple sclerosis severity (ARMSS) score. Patients in the lower 40th percentile were classified as mild, and those in the upper 40th percentile classified as severe. The 2-year follow-up data from the clinical and imaging layers were used to calculate the change from baseline, and these changes were added as new variables.

#### Multilayer network construction

Individual networks were constructed from the five layers by computing mutual information between nodes within each layer, due to the inherent nonlinear nature of biological processes. First, the networks within an individual layer were constructed, and then the networks across layers (see **Figure 2** for details on degree distribution for each layer). This step was done separately for two reasons: first to highlight the inherent differences (including biological scale) among the various layers, and second to utilize the maximum number of subjects available for each dataset. This is because not all subjects have data for both cytomics and proteomics.

Once individual layer networks were constructed, the features between layers were connected together, again with mutual information. Not all layers are interconnected, however, due to a predetermined hierarchy applied to the system (see **Figure 1g**). Ultimately, this produced a network of five connected layers, where each layer contains features from each of the five original datasets. A pipeline for the construction of the networks is shown in **Figure 1**. A second type of network was constructed using all five datasets, this time using linear correlation to define the edges, and such network was later used in the path analysis.

#### Calculation of correlation for edges

The method to calculate the edge weights in our networks was adopted from the ARACNE method (62) and simplified. The networks were constructed using mutual information, using the traditional binning method to calculate the mutual information pairwise between all the elements within individual layers and later between layers (63–65). The data for a given element are split into 10 equally spaced bins, and the probability of falling within a certain bin is calculated for each element individually as well as the joint probability for a two-point coordinate falling within a certain two-dimensional 1/10 by 1/10 size bin. The formula for the mutual information between two variables X and Y is

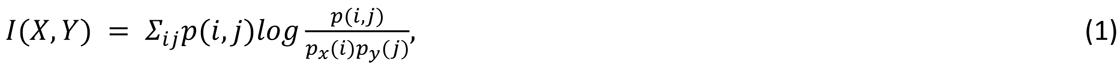

where px(i) and py(j) are the marginal probabilities for variables X and Y, respectively, and p(i,j) is the joint probability between X and Y. The python package *scikit-learn* (66) was used for the mutual information calculation.

Once the mutual information value is calculated, a threshold is needed to determine if there is indeed a correlation between the two elements. Random permutations over subjects are performed separately for both variables, and the mutual information is calculated over the permuted data. This process is repeated 1000 times, and a distribution is obtained of random mutual information values (surrogates). The mutual information value obtained from the original data is compared to the distribution of random values to determine if it is significantly higher than the distribution. The distribution is treated as Gaussian, and the original mutual information value is considered significant if it passes a z-test with p-value lower than p=0.05. Edges are placed between all significant pairs. Weights are assigned using the normalized value of mutual information, which falls between 0 (no correlation) and 1 (perfect correlation).

The combined network (later used for the path analysis) was constructed using Pearson correlation. The Pearson correlation coefficient was calculated pairwise between each of the elements included in the two datasets, using the python package *scipy* (67). An edge was defined if the p-value associated with the correlation was lower than p=0.05. Next, the value of the Pearson correlation itself was used as the weight of the edge, giving a weight that falls between - 1 (perfect negative correlation) and 1 (perfect positive correlation).

#### Path identification via Boolean modeling

The method of path identification was inspired by Domedel et al (30). The combined five-layer network was constructed using Pearson correlation, and information flow across it was analyzed using Boolean simulations. This is done to examine how perturbing the network affects nodes within the various layers, especially those representing the phenotype. The genomics network in this case was modified further, utilizing information about regulatory interactions from the Gene Regulatory Network Database (28), between the genes that are mapped to the SNPs (described further above). The exact chemical reactions between proteins and cells are ignored, giving a qualitative description of the system (29). The goal of this step is to identify differences in paths responsible for triggering immune responses in healthy subjects compared to MS patients.

For simplicity, each element in the network (from one of the five layers) is considered to be in one of two states: active/inactive. For example, this represents high/low levels of phosphorylation for proteins. The Boolean simulation begins in a random state where each element has a 50% probability of starting as active or inactive. At each step, the elements’ activation states are updated based on the sum of the states of their neighbors. The nature of the connections between elements is key, as they have either activating (positive) or inhibitory (negative) relationships. For a given node, each neighbor contributes a score based on the weight and the sign of the connection of the corresponding Pearson correlation. The total sum of the weights of the neighbors determines whether the node will be active or inactive on the next iteration.

As an example, consider the protein GSK3AB (inactive) with neighbors HSPB1 (active) and IKBA (active), as seen in **Figure 8**. Let’s say there is a positive connection between GSK3AB and HSPB1 with a weight of 0.8, and a negative connection between GSK3AB and IKBA with a weight of 0.5. Since HSPB1 is active and has a positive relationship with GSK3AB, it contributes a score of +0.8 to change GSK3AB to the active state. Since IKBA is active and has a negative relationship with GSK3AB, it contributes a score of -0.5 to GSK3AB inactive. Overall, we have a score of +0.3, so GSK3AB becomes active.

**Figure 8.**
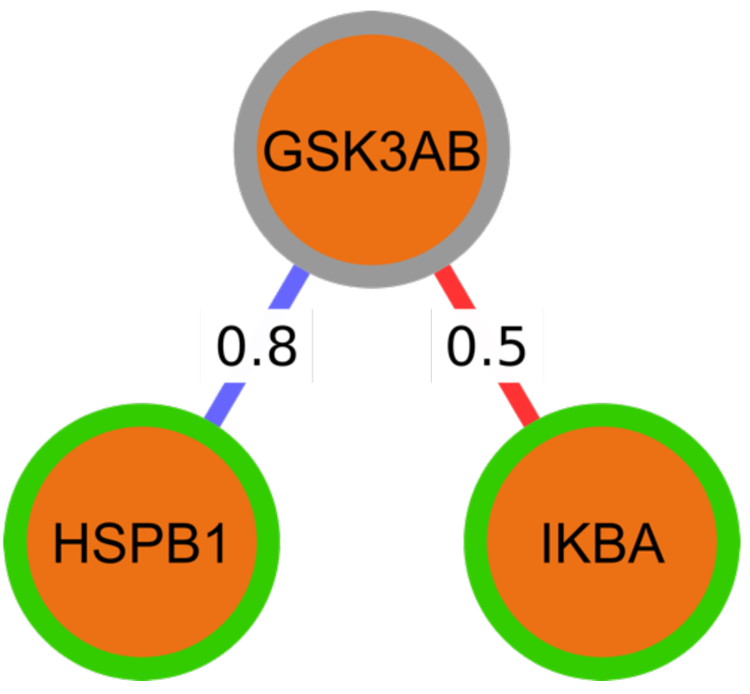
Depiction of summing weights to determine next activation state in Boolean simulations. A green border represents an active node, and a grey border represents an inactive one.

Each step of the simulation was run in this manner and continued for 100 steps. One of the MSGB scores, proteins or cells was chosen as the input, where it was manually flipped between active and inactive states with a defined period (in this case 10 iterations active, then 10 iterations inactive). This was done to examine how perturbations in the input node travel through the network and ultimately affect a given phenotype (output). The perturbations themselves represent changes between low to high values in the distribution for a given MSGB scores, protein, or cell. For the MSGB non-HLA score, the perturbations flip the value between high and low genetic risk. For a protein such as GSK3AB, the values flip between low and high phosphorylation. Finally for a cell such as B Memory, the values alternate between high and low cell counts.

Noise was also added to the system, where each element has a set probability of changing its state at each iteration. The effect of noise can is illustrated in **Figure 9**. This addition of noise reflects the inherent stochasticity in biological systems as well as prevents the simulations from simply settling directly into a fixed state. The noise was chosen to be 5% because this allows greater differences for the cross-correlation of the signals between nodes as shown in **Figure 9**. With no noise at all, many of the nodes remain either active or inactive for the majority of the simulation. This causes the cross-correlations to be too high between nodes, and the subtle differences in the strength of the connections is not seen.

**Figure 9:**
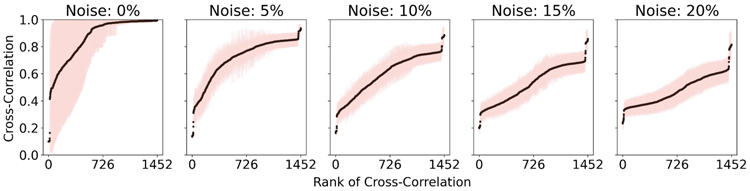
Effect of noise in Boolean simulations on the cross-correlation coefficient of the signals between nodes in the combined network. With 0% noise, a majority of the cross-correlation values are nearly 1, which does not allow the node pairs to be easily ranked based on the strength of their connections. With 5% noise, there is more deviation in the cross-correlation values, which allows the paths between a chosen source and target to be more easily identified.

**Figure 10:**
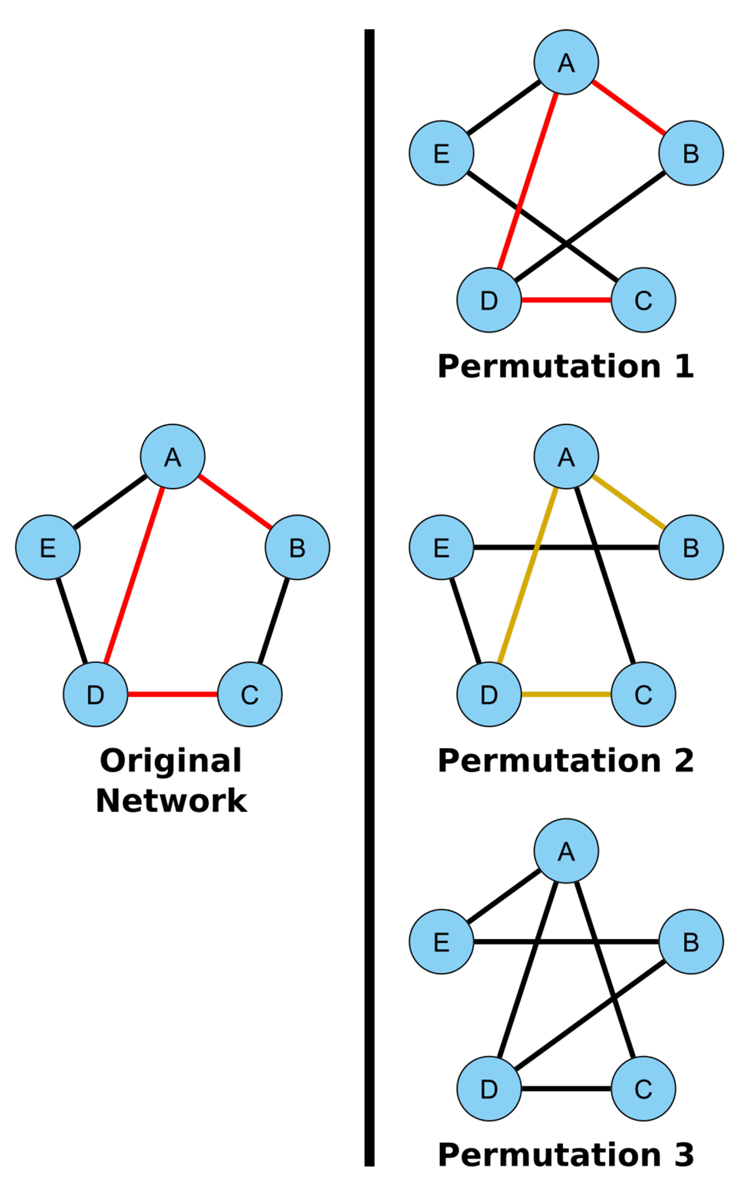
Network permutation for negative controls of paths. The five-layer network built using Pearson correlation is used as the base network. For each of the 100 repetitions, the network was permuted by swapping the edges between pairs of nodes. In permutation 1, the edge between B and C was swapped with the edge between D and E. In the permutation 2, the edge between A and E was swapped with the edge between B and C. In permutation 3, first the edge swap from the top network was applied, followed by the edge swap from the middle network. In each case, the edge swap can only be done if it does not result in two edges between the same pair of nodes. Making the permutation in this way keeps the original degree distribution of the network. The weights for each of the edges are permuted as well. This edge swapping technique is applied 10 times for each edge in the original network. After they are permuted, the top paths for each network are identified in the same manner as before. There are three possibilities for considering whether the paths from the original network appear in the paths from the permuted networks. In permutation 1, the path exists in the permuted network and furthermore was identified as a top path. In permutation 2, the original path does exist in the permuted network but was not identified as a top path. In permutation 3, the original path doesn’t exist in the permuted network at all.

Once the simulations were run, the temporal cross-correlation function was calculated between all pairs of nodes. The cross-correlation is a measure of similarity classically used in signal processing and is the same used in (30). The maximum cross-correlation (which could occur at a non-zero lag time) was determined, and its inverse is placed as a weight on the edges of the existing network, in such a way that a high correlation would correspond in this case to a low weight. In case there was no edge in the original network, no edge is defined in the new network either. A cell type or phenotype is selected as a target (output), and the most efficient paths are identified between it and the fixed source (input). An “efficient” path is defined as one in which the total sum of the weights (inverse maximum cross-correlations) of the edges connecting the source and target (called a path score) is lower than the rest. This definition favors both low number of steps and high cross-correlations between nodes within a path. A shortest path algorithm developed by (65) was used, which gives precedence to the lowest path scores.

Simulations were conducted between every possible pair of inputs (MSGB, proteins, or cells) and outputs (cells or phenotypes). Overall, the simulations aim to reveal how information flows through the entire networks, providing insight on underlying pathology in MS. This provides useful biological information, as differences in paths can be accessed between various subsets of patients (mild, severe, progressive MS, relapse-remitting MS, untreated, low-efficacy, and high-efficacy treatments). The algorithm for performing the Boolean simulations and the path identification is represented schematically in Figure 2.

In order to test the consistency of the results, we ran 100 simulations for each source, then these 100 simulations were used to calculate the cross-correlation between proteins/cells to identify the paths. We applied a jackknife resampling 10 times, first taking 90 random samples, then 80 random samples. In both cases, 9 out of 10 paths on average were identical over all protein sources and cell targets. Also, as stated in the main text, negative controls were considered by permuting the network before running the Boolean simulations. An illustration of the process for permuting the networks and identifying their corresponding paths is shown in

#### Combinatorial analysis

All possible combinations of source sources (MSGB scores, proteins, cells) and targets (cells, imaging and clinical phenotype) were used to identify top paths. The simulations were run with each protein as a source, where it remained active for 10 steps, then inactive for 10 steps. After the simulations were run for each source, and the cross-correlation values were calculated, each cell type was selected to be the endpoint for the path finding algorithm. This was performed as a screening process to create an ensemble of paths for each source/target pair. Their significance in the phenotype was assessed next.

#### Statistical analysis

The study was designed with a 1:4 ratio controls vs MS patients are based in the following reasoning: 1) the goal was the prediction of the phenotype and for such analysis only MS cases will be used; 2) controls were only used for the logistic regression comparing the diagnosis; 3) MS is heterogenous and for this reason it was expected to perform comparisons between subgroups based on disease subtype and therapy, requiring a bigger sample size for the MS group. For this reason, we designed a 4:1 ratio. Controls were collected in equal proportion from all participant centers in order to avoid center bias.

Descriptive statistics, normal distribution assessment, and class comparison analysis was performed for the five layers. The Mann-Whitney test was used due to non-normal distributions being present in both datasets. Mutual information was used in constructing the topological networks for all five layers.

#### Network statistics

Network metrics were calculated from the networks constructed using mutual information. including average degree and density. The clinical and imaging datasets lack information from healthy controls, so networks were not constructed in these cases. The average degree is given for each individual layer for healthy controls and MS patients, including those who are not treated with fingolimod (**Table 2**). Considering the omics datasets, all three of cytomics, proteomics, and genomics saw a significant increase in degree from the healthy network to all patient network at the 5% significance level. When comparing groups of patients treated with any medication versus groups excluding the patients treated with Fingolimod (a high-efficacy treatment with notable effects on cell counts in the immune system^3^, the cytomics networks saw decreases in degree in every case, and the genomics saw decreases for all patient and mild patient networks.

**Table 2.** Average degree of individual networks constructed using mutual information to define edges. These degrees do not consider the connections among layers. The superscripts (a,b) represent cases where there was a significant change when comparing degree distributions. The Mann-Whitney test was used for all pairings, due to the non-normality of the degree distributions. ^a^ Significant increase (p-val < 0.05) in degree between healthy controls and all patients. ^b^ Significant decrease (p-val < 0.05) in degree between all patients in a given subset (all, mild, or severe) and those not treated with Fingolimod.

| Average Degree |  |  |  |  |  |  |  |
| --- | --- | --- | --- | --- | --- | --- | --- |
|  | Healthy | All Patients | Patients w/o Fingo | Mild Patients | Mild w/o Fingo | Severe Patients | Severe w/o Fingo |
| **Clinical** | - | 11.2 | 11.4 | 8.0 | 8.1 | 7.8 | 7.2 |
| **Imaging** | - | 6.3 | 6.3 | 4.6 | 4.6 | 3.5 | 3.4 |
| **Cytomics** | 2.5<sup>a</sup> | 8.8 | 6.0<sup>b</sup> | 5.8 | 3.8<sup>b</sup> | 5.7 | 3.6<sup>b</sup> |
| **Proteomics** | 5.0<sup>a</sup> | 8.0 | 7.5 | 3.9 | 2.9 | 5.5 | 5.8 |
| **Genomics** | 12.2<sup>a</sup> | 13.3 | 12.6<sup>b</sup> | 12.2 | 13.0<sup>b</sup> | 12.4 | 12.3 |

## Data availability

Anonymized raw data of the Sys4MS cohort is available at MultipleMS database (www.multiplems.eu) upon reasonable request and a web interface of the networks (https://keithtopher.github.io/single_networks/<u>#/</u> and https://keithtopher.github.io/combo_networks/<u>#/</u>) and paths (https://keithtopher.github.io/fiverlayer_pathways/) are available at Github.

## Acknowledgements

We would like to thank the MS society of Norway and Italy and the GAEM foundation from Barcelona, Spain for the feedback provided to this project. We would also like to thank Dr David Gomez from the Department of Pediatric Neurology, Hospital Vall d’Hebron, Barcelona, Spain, for the review of the linear regression models. We also thank Dr. Ina Brorson for imputation of the genetic data and the research optometrists in prof. Liv Drolsums lab at Oslo University Hospital for help with the follow-up visual examinations and OCT scans and Fernanda Kropf and Ingrid Mo for technical assistance in preparation of biological samples.

**Study Funding**: This work was supported by the European Commission (ERACOSYSMED ERA-Net program, Sys4MS project, id:43), Instituto de Salud Carlos III, Spain (AC1500052); the Italian Ministry of Health (WFR-PER-2013-02361136), the German Ministry of Science (Deutsches Teilprojekt B “Förderkennzeichen: 031L0083B) and the Norwegian Research Council (project 257955). K.E.K and J.G-O. were supported by the Spanish Ministry of Science and Innovation and FEDER, under project PID2021-127311NB-I00, by the Maria de Maeztu Programme for Units of Excellence in R&D (grant CEX2018-000792-M), and by the Generalitat de Catalunya (ICREA Academia programme).

## Author Contributions

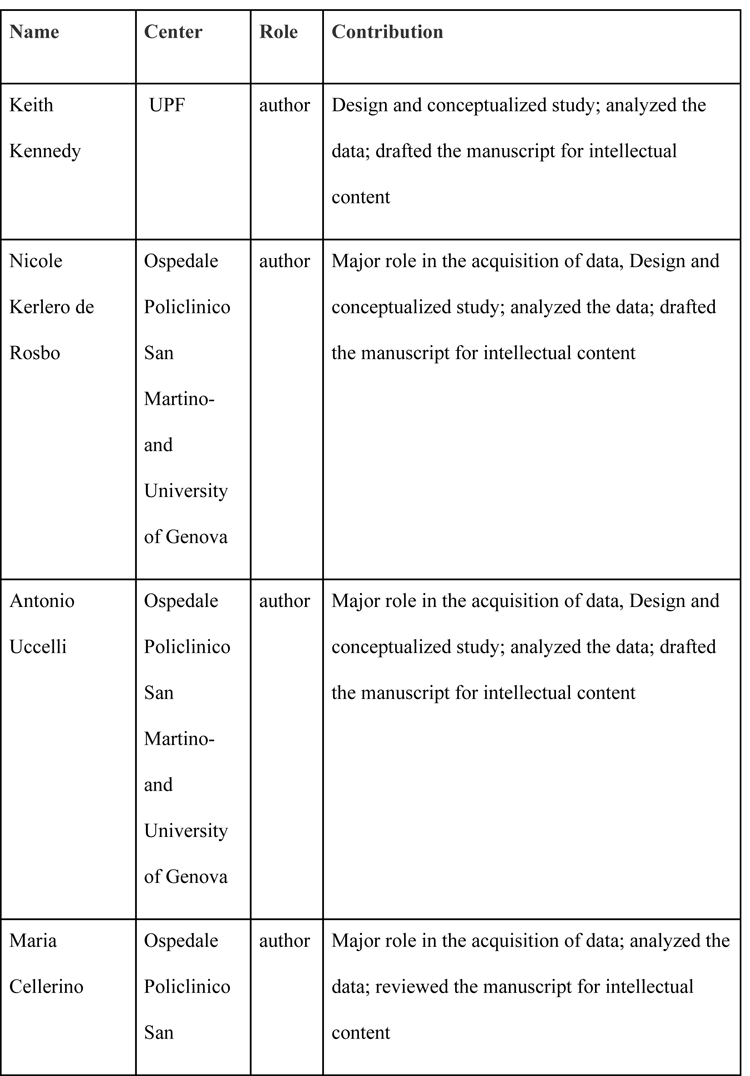
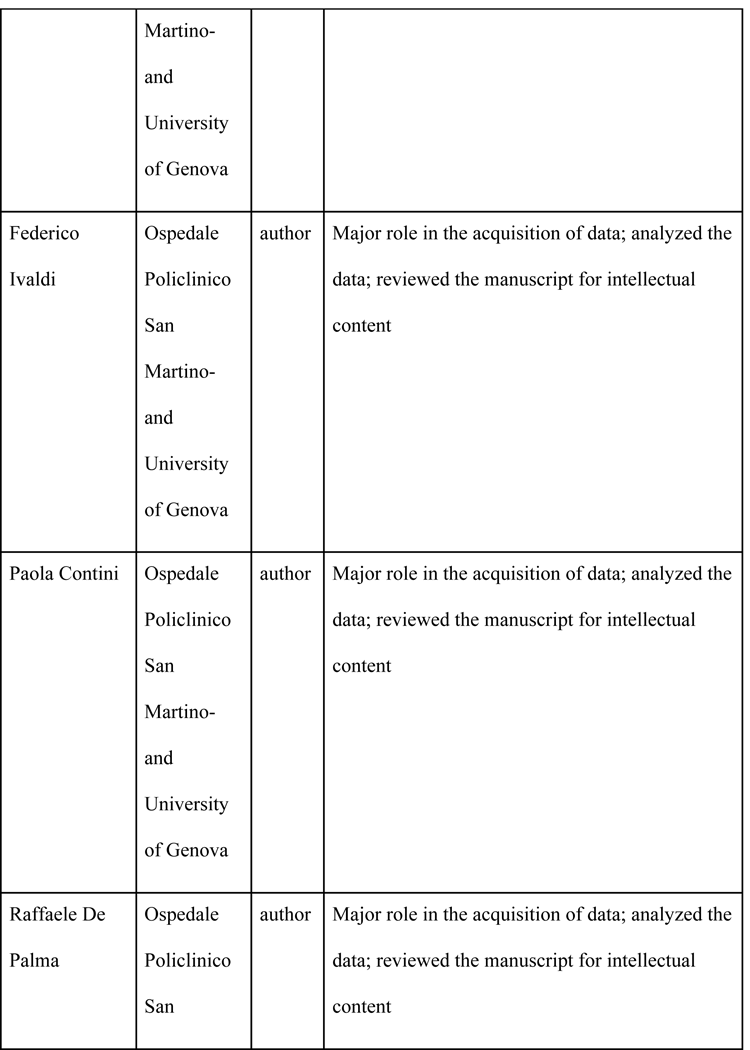
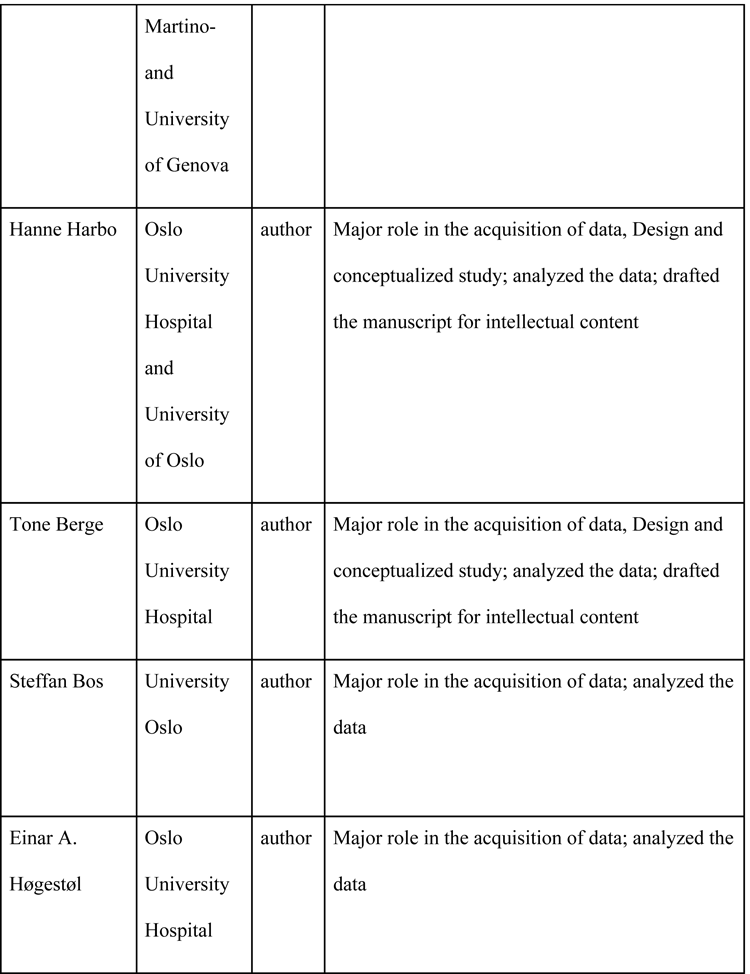
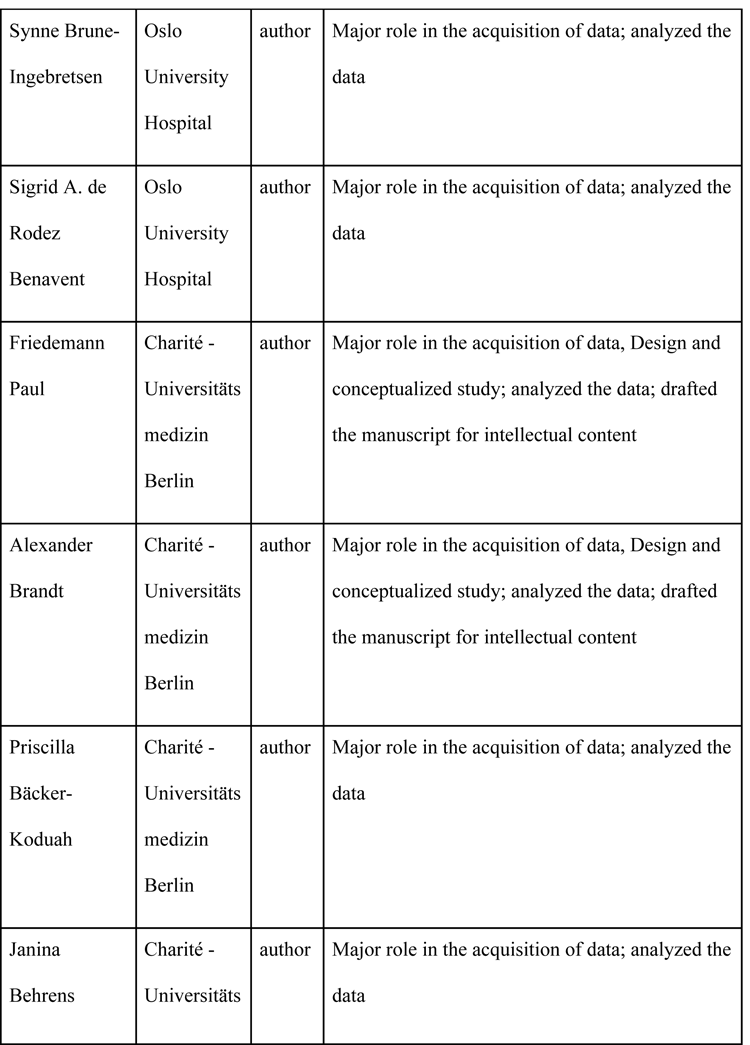
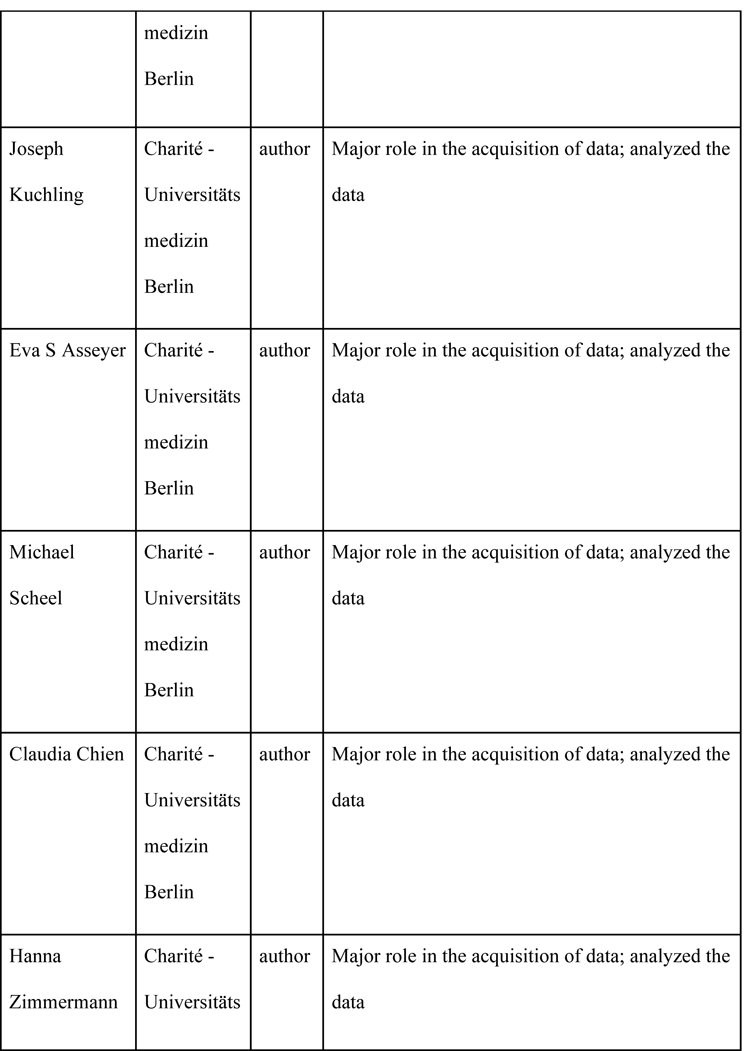
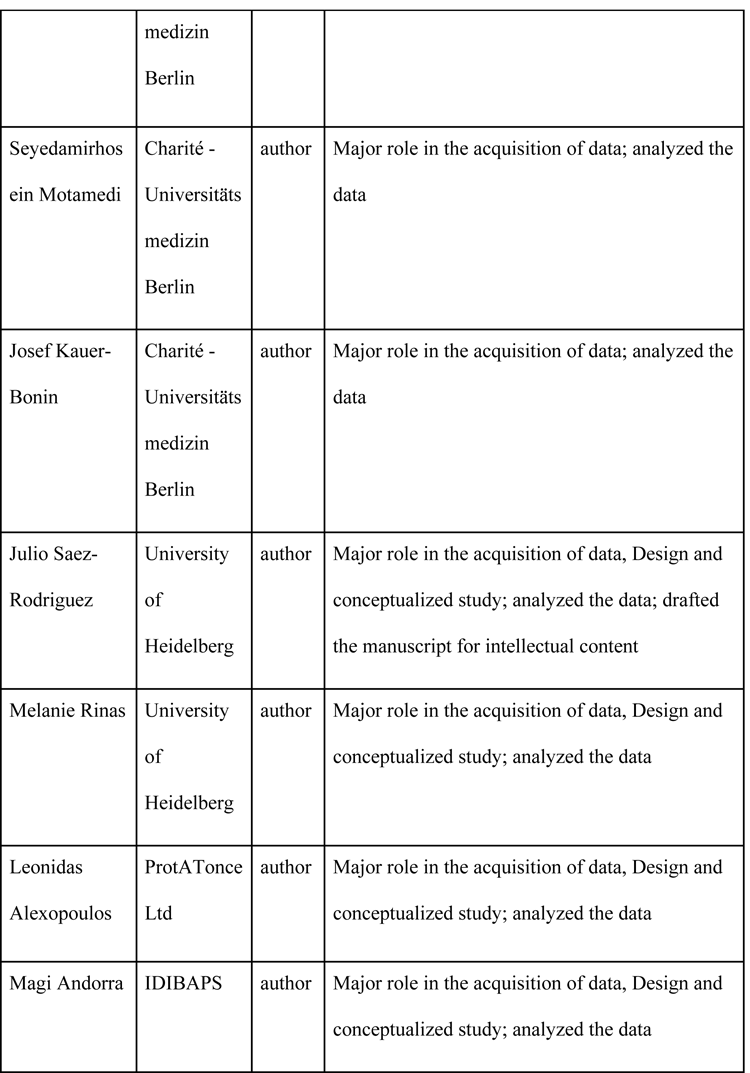
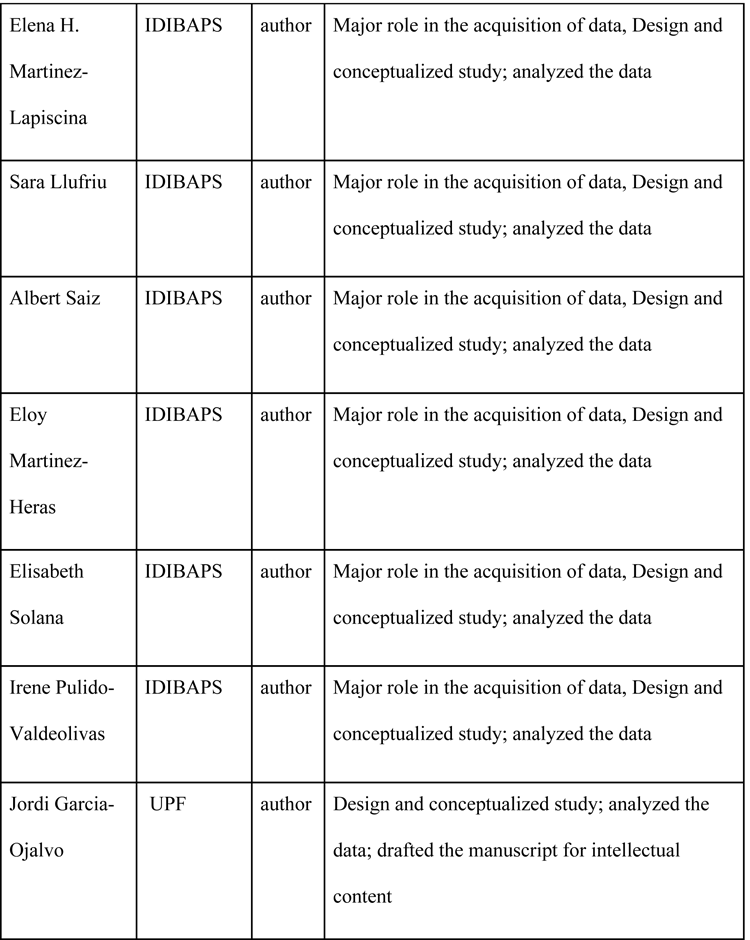
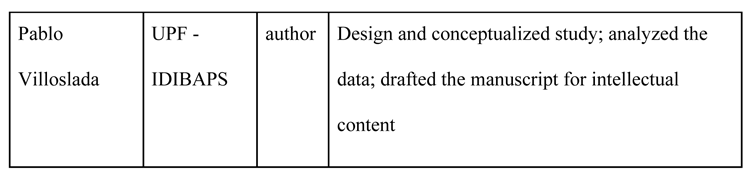

## Conflict of Interest

Keith Kennedy reports no disclosures.

Nicole Kerlero de Rosbo reports no disclosures.

Antonio Uccelli received grants and contracts from FISM, Novartis, Biogen, Merck, Fondazione

Cariplo, Italian Ministry of Health, received honoraria, or consultation fees from Biogen, Roche,

Teva, Merck, Genzyme, Novartis.

Federico Ivaldi reports no disclosures

Maria Cellerino reports no disclosures.

Hanne F. Harbo has received honoraria for lecturing or advice from Biogen, Merck, Roche,

Novartis and Sanofi.

Tone Berge has received unrestricted research grants from Biogen and Sanofi-Genzyme.

Steffan Daniel Bos reports no disclosures.

Einar Høgestøl received honoraria for lecturing and advisory board activity from Biogen, Merck and Sanofi-Genzyme and unrestricted research grant from Merck.

Synne Brune Ingebrightsen reports no disclosures.

Sigrid A. de Rodez Benavent reports no disclosures.

Friedemann Paul received honoraria and research support from Alexion, Bayer, Biogen, Chugai,

Merck Serono, Novartis, Genzyme, MedImmune, Shire, Teva, and serves on scientific advisory boards for Alexion, MedImmune, and Novartis. He has received funding from Deutsche

Forschungsgemeinschaft (DFG Exc 257), Bundesministerium für Bildung und Forschung

(Competence Network Multiple Sclerosis), Guthy Jackson Charitable Foundation, EU

Framework Program 7, National Multiple Sclerosis Society of the USA.

Alexander Ulrich Brandt is named as inventor on multiple patents and patents pending owned by

Charité - Universitätsmedizin Berlin and/or University of California Irvine for visual computing-based motor function analysis, multiple sclerosis serum biomarkers, and retinal image analysis. He is cofounder and holds shares of Motognosis GmbH and Nocturne GmbH. He serves on the executive board and is Treasurer/Secretary of IMSVISUAL. He received research support from BMWi, BMBF, NIH ICTS, the Kathleen C. Moore Foundation and the Guthy-Jackson Charitable Foundation.

Priscilla Bäcker-Koduah is funded by the DFG Excellence grant to FP (DFG exc 257) and is a Junior scholar of the Einstein Foundation.

Claudia Chien received honoraria for speaking from Bayer and research funding from Novartis, unrelated to this study.

Susanna Asseyer received a conference grant from Celgene and honoraria for speaking from Alexion, Bayer and Roche.

Janina Behrens reports no disclosures.

Julio Saez-Rodriguez declares funding from GSK & Sanofi and fees from Travere Therapeutics & Singularity Bio

Melanie Rinas reports no disclosures.

Leonidas G Alexopoulos is founder and hold stocks at ProtATonce

Magi Andorra is an employee of Hoffman-La Roche AG, yet this article is related to his activity at the Hospital Clinic of Barcelona.

Elena H Martinez-Lapiscina is an employee of the European Medicines Agency (Human Medicines) since 16 April 2019, yet this article is related to her activity at the Hospital Clinic of Barcelona and consequently, it does not in any way represent the views of the Agency or its Committees

Sara Llufriu received compensation for consulting services and speaker honoraria from Biogen Idec, Novartis, TEVA, Genzyme, Sanofi and Merck

Albert Saiz received compensation for consulting services and speaker honoraria from Bayer-Schering, Merck-Serono, Biogen-Idec, Sanofi-Aventis, TEVA, Novartis and Roche

Eloy Martinez-Heras reports no disclosures.

Elisabeth Solana received travel reimbursement from Sanofi and ECTRIMS and reports personal fees from Roche Spain.

Irene Pulido-Valdeolivas is currently an employee of UCB pharma, yet this article is related to her activity at the Hospital Clinic of Barcelona. She has received travel reimbursement from Roche Spain and Genzyme-Sanofi, European Academy of Neurology, and European Committee for Treatment and Research in Multiple Sclerosis for international and national meetings over the last 3 years; she holds a patent for an affordable eye-tracking system to measure eye movement in neurologic diseases, and she holds stock in Aura Innovative Robotics.

Jordi Garcia-Ojalvo reports no disclosures.

Pablo Villoslada has received consultancy fees and held stocks from Accure Therapeutics SL, Attune Neurosciences Inc, Spiral Therapeutics Inc, QMenta Inc, CLight Inc, NeuroPrex Inc, StimuSIL and Adhera Health Inc.

